# Human hypothalamic neural stem cells transplantation ameliorates age-associated dysfunction through SNS/eNAMPT axis activation in aged mice

**DOI:** 10.1101/2024.05.23.595504

**Authors:** Yanuar Alan Sulistio, Yuna Lee, Kelvin Pieknell, Sebin Hong, Jumi Kim, Min Jong Seok, Na-Kyung Lee, Kyu-Sang Park, Taeui Hong, Suyeon Choi, Ki Woo Kim, Dong Joo Yang, Woong-Yang Park, Kyung Yeon Han, Seul Gi Yoon, Il Yong Kim, Je Kyung Seong, Tae Yong Lee, Min Sung Kim, Min Soo Kim, Sang-Hun Lee

## Abstract

The hypothalamus is the brain region that regulates systemic body metabolism and multiple brain functions. The adult hypothalamus harbors neural stem/precursor cell (NSC)-like cells. Along with age-related body changes, the hypothalamic NSC (htNSC) population declines, indicating the potential of htNSC replacement as an anti-aging strategy. Here, we developed protocol to generate htNSCs from human pluripotent stem cells (hPSCs). Implanting the hPSC-derived htNSCs into the hypothalamus of aged mice ameliorated age-related declines in metabolic fitness, physical capacity, and cognitive function. Mechanistically, these anti-aging effects were mediated by inter-tissue communication: enhanced neuronal activity in the htNSC-transplanted hypothalamus stimulated adipose tissues to produce and release the anti-aging molecule eNAMPT into systemic circulation via the sympathetic nervous system. Concurrently, the aged inflammatory environment in the hypothalamus was alleviated by peripheral anti-aging signals. Collectively, our findings support the potential of anti- or healthy aging therapies by targeting hypothalamus.

## INTRODUCTION

Aging is a significant health concern, and developing methods to retard aging remains a prime interest of humanity. The hypothalamus, a brain region encompassing the third ventricle, functions as a high-order control center for body metabolism, energy balance, and homeostasis, all of which decline with age. Hypothalamic functions are mediated by regulating hormone production through the pituitary gland and overseeing the autonomic nervous system, thereby affecting peripheral organs like the liver, adipose tissue, skeletal muscle, bone, and intestine. Additionally, the hypothalamus has neurocircuitry connections with various brain regions, thus controlling brain functions, including circadian rhythm regulation and cognitive abilities, which also decline with age ^1–8^. Therefore, hypothalamus is integral part of inter-organ communication network that govern metabolism that can be utilized for anti-aging interventions. ^9^.

Hypothalamic functions are governed by intricate neuro-circuitry networks involving a myriad of neuronal subtypes with hypothalamus-specific neuroendocrine functions. In addition, growing evidence indicates that non-neuronal glial cells play equally important roles in the hypothalamus, such orchestrating intra-hypothalamic functions and the hypothalamic response to metabolic cues from the body, by participating in synaptic transmission, synaptic plasticity, and nutrient/hormone sensing ^10–14^. Adult hypothalamus, particularly within the third ventricle wall and median eminence, harbors cells with neural stem cell properties known as hypothalamic neural stem cells (htNSCs) or tanycytes, from which new neurons can form^15–18^. (Hereafter, ‘tanycyte’ will refer to quiescent hypothalamic NSCs in the adult hypothalamus, while ‘hypothalamic NSC’ will be used as a general term regardless of developmental and proliferation stages)^17–22^. Given that neurogenesis in the hypothalamus steeply drops after the postnatal period and is almost negligible in adulthood ^23,24^, the strategy for replenishing htNSCs might be needed to restore metabolic homeostasis during aging.

To develop a strategy for restoring metabolic balance, which is dysfunctional during aging, by targeting the hypothalamus, we differentiated hPSCs into htNSCs with neurogenic and gliogenic potential. These hPSC-derived htNSCs were then implanted into the hypothalami of aged mice. We observed improved metabolic health in the transplanted group, accompanied by several anti- aging effects. This treatment increased neuronal activity in the dorsomedial hypothalamic (DMH) region and subsided glial inflammation in the surrounding third ventricle region. Furthermore, our findings suggest that the beneficial effects in these rejuvenated mice were orchestrated by intricate inter-tissue communication between the hypothalamus and adipose tissues, mediated by the sympathetic nervous system

## RESULTS

### Generation of htNSCs from hPSC-derived hypothalamus-like organoids

Despite the great clinical importance of the hypothalamus, studies on hypothalamic functions have been made exclusively in animal models. Their translation to humans remains nascent, mainly due to the inaccessibility of human hypothalamic tissues and cells. Several studies have generated hypothalamic neuronal cell types from human pluripotent stem cells (hPSCs) ^25–27^. Given the functional significance of glia and htNSCs in the hypothalamus, however, it is compelling to develop research platforms that can provide diverse hypothalamic cell types, including non-neuronal cells, of human origin. We aimed to develop a method to differentiate hPSCs into htNSCs with neuro- and gliogenic potential.

In this work, we leveraged the principles and advantages of the organoid-based method to prepare human htNSCs from hPSCs ^28^. To this end, human embryonic stem cells (hESC, H9) were induced to differentiate into hypothalamus-like organoids in the presence of small molecules for inducing hypothalamic fate ^25–27^ (Fig. 1A and Suppl. Fig. S1A-B). The combinations, concentrations, and treatment periods of the hypothalamic patterning molecules were optimized based on high expression of the hypothalamus–specific markers *RAX*, *NKX2-1*, *OTP*, *ISL1*, and *SF-1* and low expression of the midbrain marker Engrailed-1 (*EN1)* (Suppl. Fig. S1A). Treatment with SHH and purmorphamine during early organoid differentiation was indispensable for hypothalamus-specific marker expression (Fig. 1B and Suppl. Fig. S1A). Further addition of CHIR, CNTF, IWP-2, XAV, and FGF8, which have been reported as molecules for hypothalamic patterning ^25–27^, did not enhance but lower hypothalamic marker expression (Suppl. Fig. S1A). Cells dissociated from the organoids on DIV 20 were efficiently sub-cultured as monolayer. Generated htNSCs expressed markers specific for general NSCs (SOX2, NESTIN), proliferating cells (KI67), and hypothalamic specific markers, indicating efficient generation of a homogenous htNSC culture from the early stage of the organoids (Fig. 1B‒C). The expressions of hypothalamus-specific markers and the percentage of Nestin+ and RAX+ htNSCs in our method were significantly higher compared to those observed in htNSC cultures generated using other published protocols ^25,26^ (Suppl. Fig. S1B-E). To further confirm their hypothalamic identity, we performed RNA sequencing (RNA-seq) on our cultured htNSCs. When comparing our htNSC transcriptome profile to various adult human brain transcriptome (GTEx Database, ages of 22-68 years), the htNSC was clustered together with the human adult hypothalamus, signifying a high hypothalamic regional identity (Fig. 1D).

**Fig. 1.**
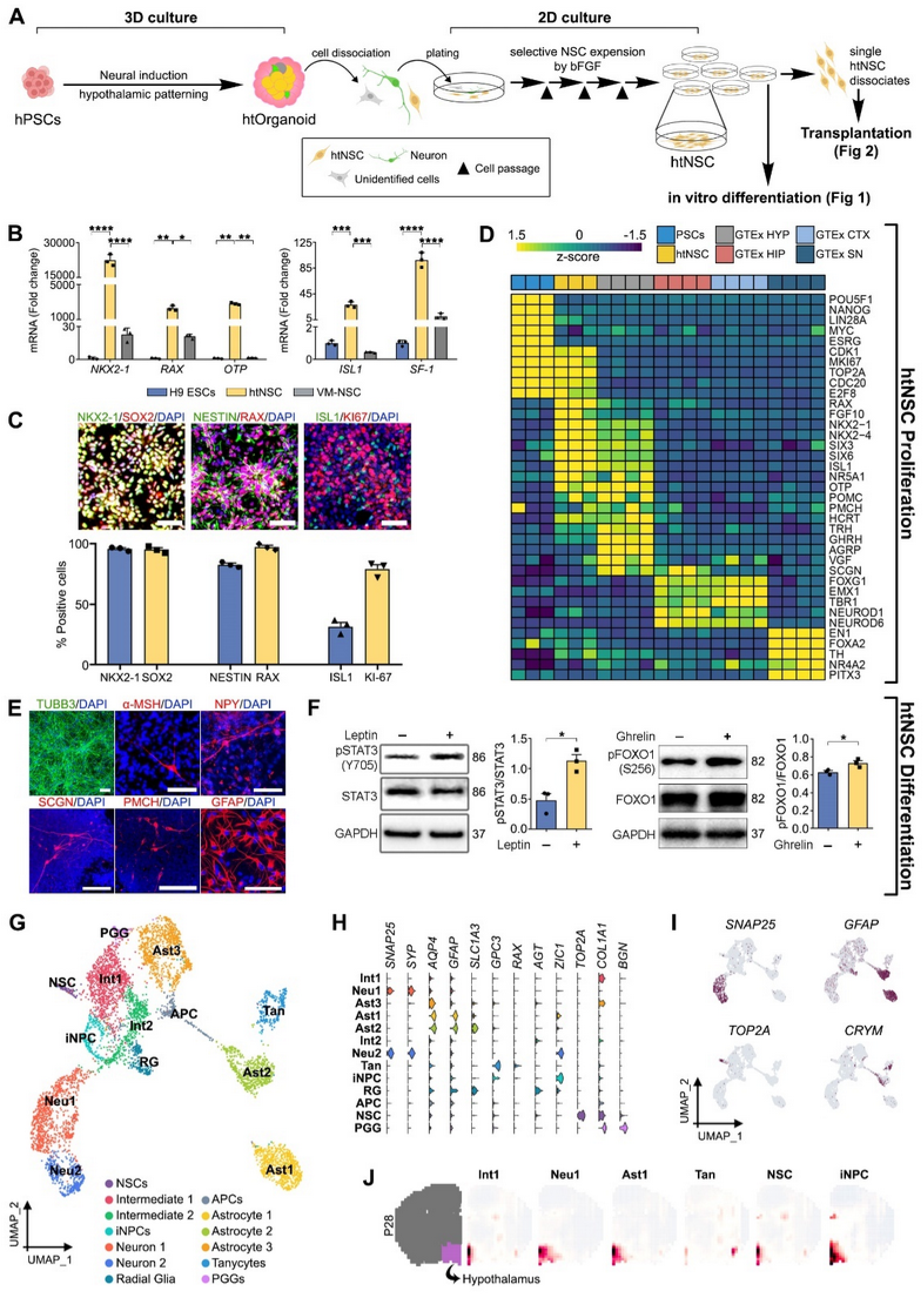
Derivation of hypothalamic neural stem cells (htNSCs) from hPSC-derived hypothalamus-like organoids. **A.** An overview of the protocol to yield htNSCs with proliferative and neuro-/gliogenic potential from hPSC-derived organoid cultures. **B.** mRNA expression of htNSC-specific marker genes (*NKX2-1*, *RAX, OTP*, *ISL1*, *SF1*) assessed by qRT-PCR. The expression levels in the H9 hESC-derived htNSCs were compared with those in undifferentiated hESCs and hESC-derived ventral midbrain-type neural stem cells (VM-NSCs). n=3.^a,b^ **C.** Immunocytochemical detection of generic NSC (SOX2, NESTIN, KI67) and htNSC-specific markers (NKX2-1, RAX, ISL1). Scale bars, 100 µm. n=3.^a^ **D.** Heatmap comparison of our H9 hESC-derived htNSC transcriptome with various human adult brain regions in the GTEx database. HYP: hypothalamus; HIP: hippocampus; CTX: cortex; SN: substantia nigra. **E.** Terminal differentiation of htNSCs to yield hypothalamus-specific neuronal and astroglial progeny. The cellular identities were evaluated on htNSC differentiation day 40 by the expression of general neuronal (TUBB3, MAP2, RBFOX3, SYN1), hypothalamus neuron subtype (α-MSH, NPY, SCGN, PMCH), and astroglial (GFAP, VIM, AQ4, A2B5) markers. Scale bars, 100 µm. **F.** Cells differentiated from htNSCs acquired the hypothalamus-specific functions of responding to leptin and ghrelin ligands. Intracellular STAT3 and FOXO1 activation were evaluated in the differentiated htNSC cultures before and after exogenous leptin and ghrelin treatments, respectively. n=3.^a,b^ **G.** Uniform manifold approximation and projection (UMAP) plot of 6,394 single cells projected into 13 cell clusters (indicated by color). NSCs, neural stem/progenitor cells; Int1, intermediate progenitor cells 1, Int2, intermediate progenitor cells 2; iNPCs, interneuron progenitor cells; APCs, astrocyte progenitor cells; Neu1, neurons 1; Neu2, neurons 2; Ast1, astrocytes 1; Ast2, astrocytes 2; Ast3, astrocytes 3; RG, radial glia; PGG, proteoglycan-secreting glia; Tan, tanycytes. **H–I.** Violin (B) and feature (C) plots of representative cell-specific genes from the differentiated cell clusters. **J.** Hypothalamic brain identity of the cell clusters validated by voxel-based spatial brain mapping onto the mouse brain at postnatal day 28 (from the Allen Brain Institute). ^a^Data shown are means ± SEM. ^b^*p < 0.05; **p < 0.01; ***p < 0.001; ****p < 0.0001 by two-tailed Student’s t testing.

Additionally, the htNSCs were expandable for extended periods (Suppl. Fig. S2A), could be stored and re-cultured from liquid N2 without loss of viability or naïve htNSC-specific marker expression (Suppl. Fig. S2B), establishing the htNSC culture derived from hPSC-organoids as a stable and readily available source of human htNSCs. Proliferation, indicated by PDL, decreased in late passages (Suppl. Fig. S2A), signifying replicative senescence.

### Differentiation phenotypes of the hPSC-derived htNSCs

Upon terminal differentiation, the htNSCs could differentiate into mature neurons expressing (TUBB3). It also expressed hypothalamus-specific neuropeptides such as POMC, NPY, SCGN, and PMCH (Fig. 1E). Astrocyte was detected in the differentiated htNSC cultures (Fig. 1E), which increase in proportion in the later passages (Suppl. Fig. S2C). Hypothalamic neurons and astrocytes sense the nutritional status of the body through peptides such as leptin and ghrelin that are generated in peripheral organs. An administration of leptin to the differentiated htNSC cultures activated the major leptin-mediated STAT intracellular pathway, which was shown by increased phosphorylated STAT3 (pSTAT3) levels in Western-blot analysis (Fig. 1F, left). In addition, the phosphorylation of FOXO1, the intracellular signal transducer of ghrelin, occurred when the differentiated htNSC cultures were treated with ghrelin (Fig. 1F, right) ^29^ indicating that the differentiated hypothalamic cells functionally sensed nutritional cues. Using this protocol, htNSCs with neurogenic and gliogenic capacities were also generated from the human induced pluripotent stem cell (hiPSC) lines CMC-hiPSC-011 and CS-001(Suppl. Fig. S3).

Next, we performed a single-cell RNA-seq (scRNA-seq) analysis of the differentiated htNSC culture after seventy days of terminal differentiation. After filtering doublets and clusters cells with low-quality reads, we used 6,394 cells for the analysis. Unsupervised clustering of the dataset revealed thirteen clusters, which were projected onto a uniform manifold approximation (UMAP) map for visualization. On the basis of the expression levels of well-characterized marker genes ^30–32^ alongside with automated identification using scType, the cluster identities were designated as proliferating neural stem cells (NSCs), intermediate progenitor cells (Int1, 2), interneuron progenitor cells (iNPCs), astrocyte progenitor cells (APCs), neurons (Neu1, 2), astrocytes (Ast1, 2, 3), radial glia (RGs), proteoglycan-secreting glia (PGGs), and tanycytes (Tan) (Fig. 1G-I and Suppl. Fig. S4A). For unbiased confirmation of the hypothalamic fate, we mapped the gene expression signatures of the cell clusters to the mouse brain transcriptome (at post-natal day 28) in the Allen Developing Brain Atlas using Voxhunt ^33^. Significantly, our scRNA-seq profiles showed that every identified cluster in the differentiated htNSC culture precisely corresponded to the hypothalamic region of the atlas (Fig. 1J and Suppl. Fig. S4B). Furthermore, the progenitor and intermediate cell clusters (Int1, Int2, iNPC, APC, NSC) were enriched to the hypothalamic region adjacent to the ventricular area, where adult neurogenesis occurs (Fig. 1J and Suppl. Fig. S4B). To provide additional confirmation, we cross-referenced our single-cell transcriptome data with published datasets of human fetal hypothalamus (gestational week (GW) 18-22) (Suppl. Fig. S4C-D) ^34^. The clustering analysis indicates that our single-cell transcriptome data align closely with the human fetal hypothalamus datasets (Suppl. Fig. S4E-F). Expectedly, oligodendrocytes and microglia were absent in the differentiated htNSC (Suppl. Fig. S4G). Next, we performed a gene regulatory network analysis using pySCENIC and identified the potential upstream master regulator (regulon) of each cell cluster (Suppl. Fig.S4H).

The hypothalamus is a complex brain area that contains myriad subtypes of neurons with neuroendocrine and neurotransmitter identities. To unravel the identity of the neuronal populations in our differentiated htNSC cultures, we performed further clustering on the neuronal clusters (Neu1 and Neu2). Upon re-clustering based on the signature transcripts, we detected 8 subclusters of neurons (n1–n8) (Suppl. Fig. S5A), comprising neuronal clusters with different neurotransmitter identities, such as glutamatergic (n6), GABAergic (n2), dopaminergic (n8), and dopaminergic/glutamatergic (n1, n3, n4), and clusters with specific hypothalamic neuroendocrine identities, such as those expressing *SCGN* (n5), *PMCH* (n4), *TRH* (n3), and *NR3C1* (n1) (Suppl. Fig. S5B-C). In addition, the neuronal subclusters were classified by their unique transcriptional factors and regulons (Suppl. Fig. S5B-C). Of note, *VGF*, a gene that is highly enriched in the adult hypothalamic neurons ^35,36^, was widely expressed throughout the neuronal clusters (Suppl. Fig. S5B).

The APC cluster branched into three astrocyte trajectories (Suppl. Fig. S5D). Astrocyte cluster Ast2 is regarded as a homeostatic astrocytic population that expresses *SLC1A3* (GLAST), a protein that provides metabolic support for neighboring neurons (Suppl. Fig. S6F). Additionally, Ast2 could be involved in neuroendocrine functions because of its specific expression of genes such as *SLCO1C1* (thyroid hormone transporter), *CRYM* (Crystalin Mu, a thyroid hormone binding protein), and *SCG2* (secretogranin II, a neuroendocrine secretory protein) (Suppl. Fig. S5E). By contrast, Ast1 preferably expressed genes related to immune function such as *RSAD2* and *TRIL* (Suppl. Fig. S5E). Moreover, interferon-induced genes, such as *ISG5*, *IFIT*, and *STAT1*, were also specifically expressed in Ast1, suggesting an immunogenic phenotype. Aside from the “classical” homeostatic and reactive astrocytes, we also detected a third type of astrocyte (Ast3) that was derived from common progenitor cells (APCs) and expressed astrocyte-related transcription factors (*SOX9*, *ID3*, *BHLHE41*) and common astrocyte markers (*APOE*, *AQP4*, *SPARCL1*, *SPARC*) but was devoid of *GFAP* (Suppl. Fig. S5E). Ast3 is characterized by gene expression specific for not only tight junction formation (*CLDN5*, *ALCAM*, *COL1A2*), but also the astrocyte population in the adult subventricular zone (*Meis2*, *Col1a2*), which interacts with ependymal cells or pericytes ^37–39^. Of note, a cluster was designated as a tanycyte-like population because it exclusively expressed tanycyte-specific markers such as *RAX*, *FGF10*, *COL25A1*, *GPC3*, and *DIO2* (Suppl. Fig. S5E). Interestingly, *LEPR* (leptin receptor) expression was detected in the tanycyte cluster (Suppl. Fig. S5E), supporting the involvement of tanycytes in metabolic homeostasis via leptin signaling ^19^. The presence of tanycyte-like cells post-differentiation, further suggest hypothalamic identity of our htNSC.

### *In vitro* htNSC differentiation recapitulated hypothalamic brain development

Next, we used Monocle 3 to perform pseudo-temporal ordering of the transcriptional dynamics of the cell clusters that were differentiated from human htNSCs. Consistent with brain development, in which neurons and astrocytes arise sequentially from common progenitors, the UMAP clustering shows that the intermediate progenitor cell clusters (Int1, 2) were bifurcated into neurons (Neu1, 2) and the APC cluster that gave rise to the astrocytic populations (Ast1–3) (Suppl. Fig. S6A). Several genes showed temporal dynamic changes in gene expression that recapitulated those in the developing brain. For example, a steep decrease in the expression of the proliferation marker *CDK1* at an early time point was followed by a surge in the expression of the pro-neural gene *ASCL1* at the late progenitor cell stage and increased expression of the neuron-specific *TUBB3* and *SNAP25* at the terminal stage of the neuronal trajectory (Suppl. Fig. S6B). The *SLIT-ROBO* signal has recently been identified as a critical regulator of hypothalamus-specific brain development ^40^. Of note, the *SLIT-ROBO* signal genes in the differentiated htNSC cultures showed pseudo-time patterns similar to those of *in vivo* hypothalamic brain development (Suppl. Fig. S6C), indicating our *in vitro* htNSC differentiation recapitulates hypothalamus development *in vivo*.

Thus far, the developmental origin of tanycytes has been unidentified. Interestingly, consistent with a developmental trajectory analysis suggesting that tanycytes originate from the astrocyte population during hypothalamic brain development ^40^, our pseudo-time analysis indicates that tanycyte-like cells (Tan) developed from the early Ast2 population (Suppl. Fig. S6D-E).

### Age-associated declines in physical and cognitive activity were ameliorated in mice that received hPSC-derived htNSCs

Physical strength, muscle endurance, motor coordination, memory, and cognitive function are all declining with age ^41^, and those age-associated changes are linked to hypothalamic dysfunction in regulating body metabolism and homeostasis ^42^. A previous study has shown that age-associated impairment of physical and cognitive activity could be delayed by implantation of htNSCs derived from mouse pups ^10^. To take a critical step toward the clinical translation of the htNSC-based anti-aging strategy, we transplanted single dissociates of human htNSCs (passage 2-3) prepared from the hESC-derived hypothalamus-like organoids into the hypothalami of mice at 12-months of age (Fig. 2A). The mouse htNSCs derived from newborn mice, which were used in the previous study ^10^, survived poorly after implantation into the mouse hypothalamic region, and thus genetic engineering with dominant recombinant IκB to inhibit the NF-κB inflammatory signal was needed to overcome the survival problem ^10^. By contrast, upon transplantation into the medio-basal hypothalamus (MBH), the hPSC-derived human htNSCs survived and integrated efficiently into the host hypothalamus, forming human-specific marker (hNCAM or human mitochondrial-specific antigen)-positive grafts, portions of which expressed undifferentiated htNSC (RAX), differentiated neuronal (NeuN), and astrocytic (human GFAP, hGFAP) markers 2–4 weeks post-transplantation (Fig. 2B–C).

**Fig. 2.**
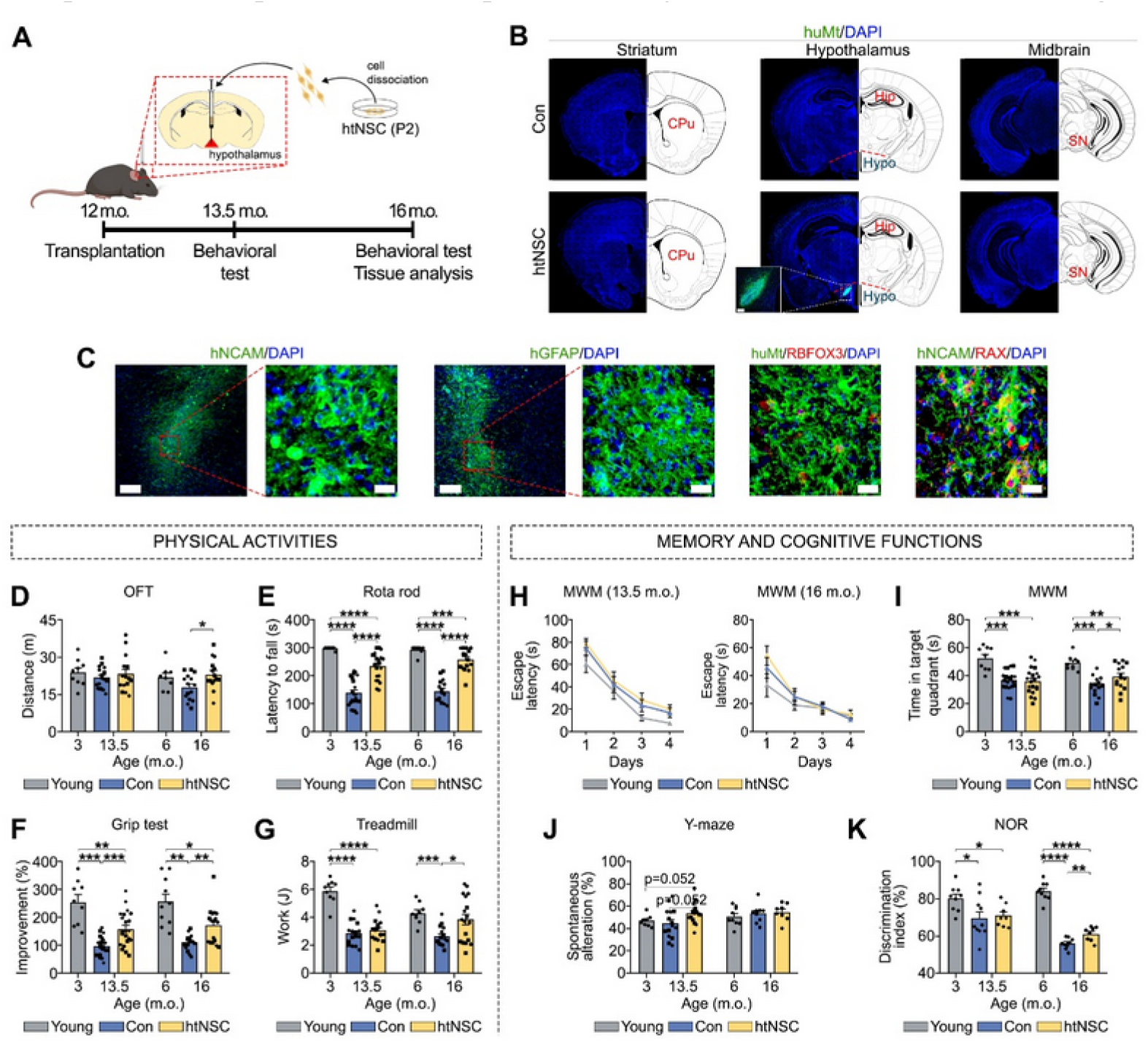
Age-associated decline in physical and cognitive activities is rescued by intra-hypothalamic htNSC transplantation. **A.** Experimental scheme for the animal study. hESC-derived htNSCs (P2) were dissociated into single cells and transplanted into the medio-basal hypothalami of 12-month-old male mice. Behavioral tests were performed 1.5 months and/or 4 months post-transplantation, and tissue analyses were conducted 4 months post-surgery. Four rounds of transplantation experiments were conducted using mice and donor human htNSCs that were independently prepared. **B.** Overview images depict the human htNSC-derived graft localized in the mouse hypothalamus but not in other brain regions. The htNSC-derived graft was identified using a human mitochondria-specific antibody (huMt). CPu, caudate putamen; Hip, hippocampus; Hypo, hypothalamus; SN, substansia nigra. **C.** Identification of cell types derived from the grafted htNSCs using antibodies specific for human-species (hNCAM, huMt), human-specific astrocytes (hGFAP), htNSCs (RAX), or neurons (RBFOX3) at 2–4 weeks post-transplantation. Scale bars, 50 µm or 200 µm. **D–G.** Physical activity, muscle coordination, strength, and endurance were estimated using the open filed test (OFT, C), rotarod (D), grip (E), and treadmill performance (F) tests. n= 9-10 (young), 15-22 (control), 15-22 (transplanted).^a,b^ **H–K.** Memory and cognitive functions were evaluated using the Morris water maze (MWM, G,H), Y-maze (I), and novel object recognition (NOR, J) tests 4 months post-surgery. n= 9 (young), 9-20 (control), 8-20 (transplanted).^a,b^ ^a^Data shown are means ± SEM. The data presented in D-K represent the combined data from at least two independent datasets. ^b^*p < 0.05; **p < 0.01; ****p < 0.0001 by one-way Welch analysis of variance.

To ensure reproducibility, we designed multiple cohorts of cell transplantation with independently prepared human htNSCs and aged mice. In those consecutive trials of htNSC transplantation, the mice that received transplantation displayed a significant improvement in exploratory behavior and locomotor activity (Fig. 2D), coordination (rotarod) (Fig. 2E), muscle endurance (grip test) (Fig. 2F), and treadmill performance (Fig. 2G), compared with the age-matched sham-operated control mice (control, culture media-injected). Furthermore, the long-term and spatial memory and cognitive functions of the grafted mice were significantly improved in the Morris water-maze (MWM; Fig. 2H–I) and novel object recognition tests (Fig. 2J). The physical activities and memory/cognitive functions in the grafted old mice were also compared with the young mice (aged 3-6 months) (Fig. 2 and Suppl. Fig. S7).

### Improvement of body energy metabolism and tissue aging by intra-hypothalamic htNSC transplantation

The grafted mice were leaner, devoid of age-dependent body weight gain during the post-transplantation period (12–16 months) that was manifested in the vehicle-injected control mice (Fig. 3A). There was no difference in the food intake and physical activity changes only occurred during the dark phase (Fig. 3B–C). Consequently, the htNSC-grafted mice displayed greater energy expenditure in the metabolic cage compared to control aged mice (Fig. 3D). Body composition analysis indicates that the htNSC-grafted mice had markedly lower fat mass with increased lean mass indicating lower adiposity (Fig. 3E). Accordingly, lipid droplet size was greatly reduced in the inguinal white adipose tissue (iWAT), epididymal white adipose tissue (eWAT), and intrascapular brown adipose tissue (iBAT) of the grafted mice than the control mice (Fig. 3F–H). Similarly, hepatic steatosis in the aged mice was markedly decreased in the transplantation group (Fig. 3I).

**Fig. 3.**
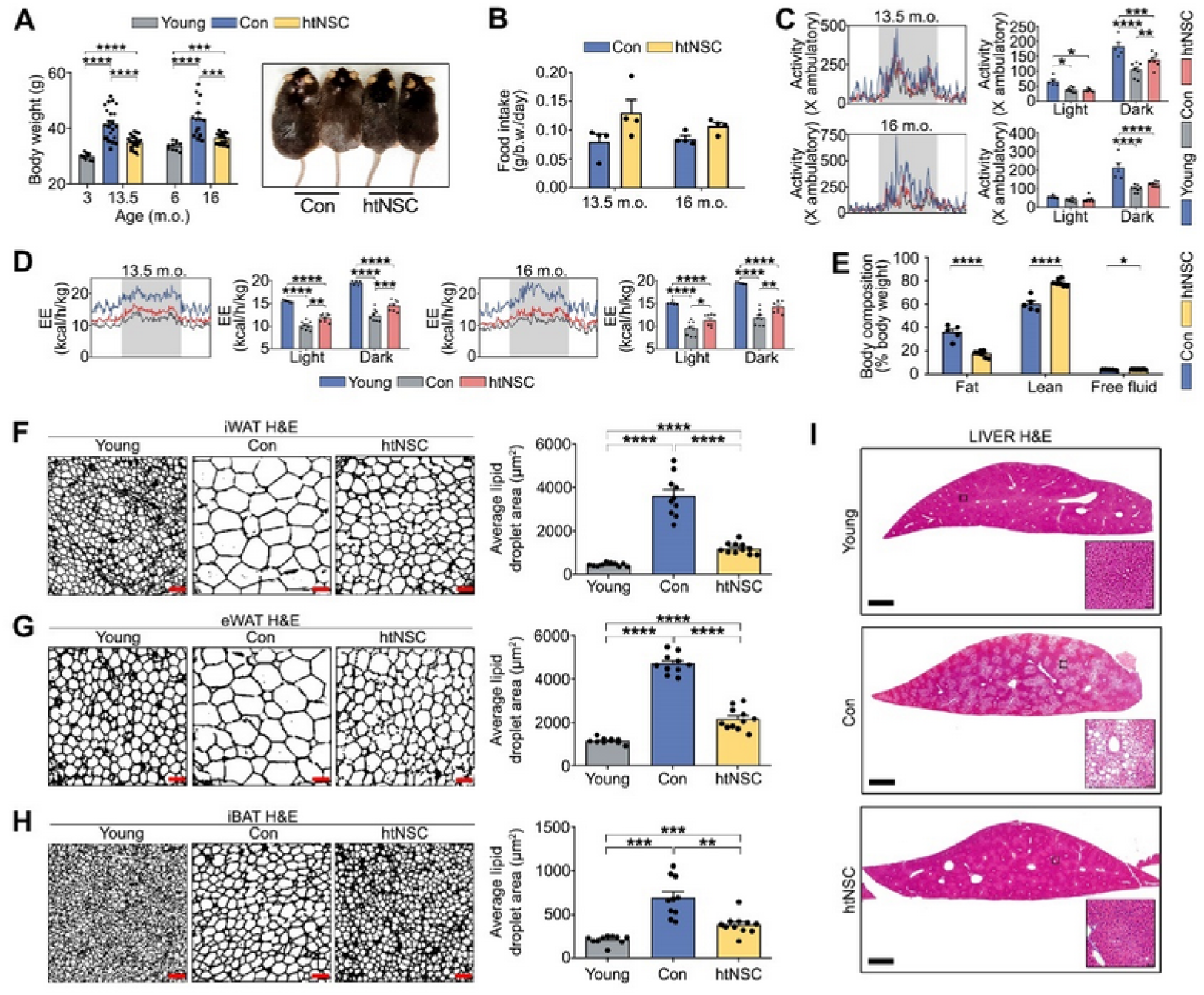
Intra-hypothalamic human htNSC transplantation improved body energy metabolism and tissue aging in aged mice. **A.** The body weights of the control (vehicle-injected) and transplanted mice 1.5 months and 4 months post-transplantation (left) and representative photographs of the mice at 16 months old (right). n= 10 (young), 15 (control), 17 (transplanted).^a,b^ **B.** Daily food intake (normal chow diet) of the control and transplanted mice. n= 5 (control), 7 (transplanted).^a,b^ **C.** Measurement of x-ambulatory activity during the light and dark phases of the day at 1.5 months post-transplantation. n= 5 (young), 8 (control), 8 (transplanted).^a,b^ **D.** Energy expenditure measured in metabolic cages. n= 5 (young), 8 (control), 8 (transplanted).^a,b^ **E.** NMR-based measurement of the fat and lean masses and free fluid volumes in the vehicle-treated control (n= 5) and transplanted (n = 7) mice.^a,b^ **F–H.** Quantified lipid droplet areas (right) with representative H&E-stained tissue section images (left) of the iWAT (F), eWAT (G), and iBAT (H) liver from the young (n = 10), control (n= 10) and transplanted (n = 11) mice. Scale bars, 50 µm. ^a,b^ **I.** Representative H&E-stained images of liver from the young (n = 10), control (n= 10) and transplanted (n = 11) mice. Scale bars, 1 mm. ^a^Data shown are means ± SEM. The data presented in A, C-D, and F-I represent the combined data from at least two independent datasets. ^b^*p < 0.05; **p < 0.01; ****p < 0.0001 by one-way analysis of variance or two-tailed Student’s t testing.

Growth/differentiation factor 15 (GDF15) and fibroblast growth factor 21 (FGF21) are key secretory proteins that stimulate peripheral tissues to increase energy expenditure and lipid consumption ^43–46^. Hepatic GDF15 and FGF21 proteins and their upstream regulator ATF4 were significantly upregulated in the livers of the grafted mice (Fig. 4A). Additionally, FGF21 was also increased in iWAT and eWAT of the grafted mice, which was accompanied by the upregulated protein expression of MT-CO1 and SDHB, mitochondrial proteins that execute energy consumption/thermogenesis (Fig. 4B–C) ^47,48^. Consistently, rectal temperature was significantly higher in the htNSC-grafted mice than the controls (Fig. 4D). Interestingly, this metabolic signature closely resembles that observed in leucine-deprived diet, which is known to promote metabolic health and prolong lifespan ^49^. The hypothalamus influences the metabolism and physiology of adipose tissues through the sympathetic nervous system (SNS) ^50^. Messenger RNA expression of β1, 2, and 3 adrenoreceptors (*Adrb1*, *Adrb2*, and *Adrb3*) and lipolytic genes (*Dio2* and *Lipe*) was significantly upregulated in the eWAT of the grafted mice (Fig. 4E), implying SNS activation in the WAT of the grafted mice. Along with efficient energy metabolism, the mice that received intra-hypothalamic htNSC transplantation showed significantly better glucose tolerance than the control mice (Fig. 4F). A bone µCT examination showed osteoporotic changes in the tibias and femurs of the control mice at age 16 months (Fig. 4G-H). That bone loss was greatly prevented by intrahypothalamic htNSC transplantation, as evidenced by a significant increase in bone mass density (BMD), bone volume/total volume (BV/TV) ratio, trabecular thickness (Tb.Th.), and trabecular number (Tb.N.) in the grafted mice (Fig. 4G-H). Notably, the increase in bone density coincided with a substantial decrease in the osteoclast population among the transplanted mouse group (Fig. 4I), suggesting that suppression of bone resorption underlies the observed enhancement in bone density.

**Fig. 4.**
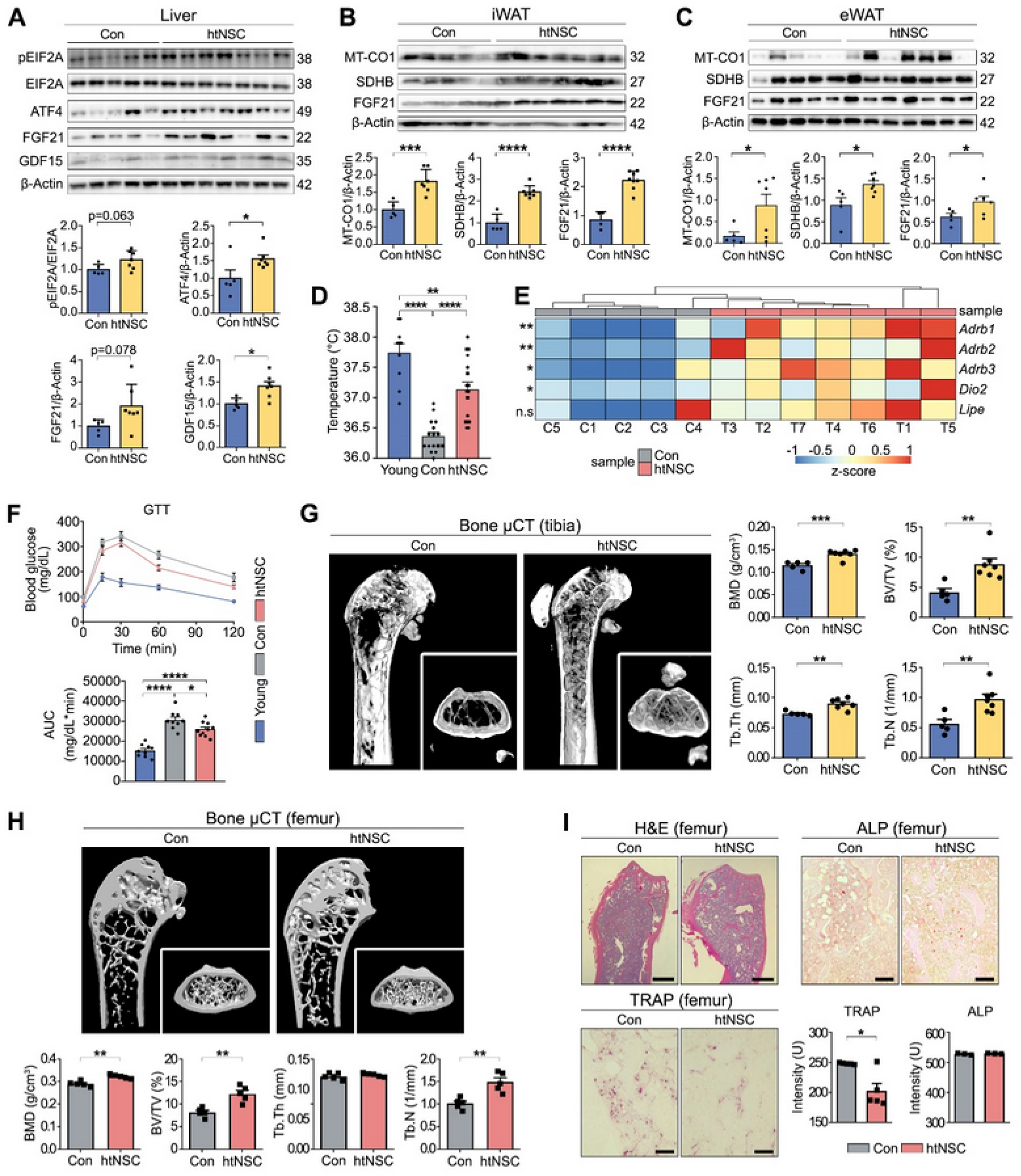
Body energy and bone metabolism changes ameliorated in aged mice that received htNSC grafts. **A–C.** Immunoblot-based quantification of the key regulator proteins for energy metabolism and thermogenesis (browning) in the liver (A), iWAT (B), and eWAT (C) of the control (n=5) and transplanted (n=7) mice. ^a,b^ **D.** Rectal temperature. n= 10 (young), 16 (control), 17 (transplanted).^a,b^ **E.** qPCR-based quantification of gene expression for β1, 2, and 3 adrenoreceptors *(Adrb1*, *Adrb2*, and *Adrb3*), and lipolytic genes (Dio2 and Lipe). n= 5 (control), 7 (transplanted).^a,b^ **F.** Glucose tolerance test. Data represent blood glucose levels over the time points after glucose injection (2 g/kg, i.p.) (left) and the integral of the glucose levels during the post-injection period (right). n= 9 (young), 10 (control), 10 (transplanted).^a,b^ **G.** Representative μCT bone images of tibia (left) and the indices of BMD, BV/TV, Tb.Th., and Tb.N. (right), which increased significantly in the transplanted mice. n= 5 (control), 7 (transplanted).^a,b^ **H.** Representative μCT bone images of femur (top) and the indices of BMD, BV/TV, Tb.Th., and Tb.N. (below), which increased significantly in the transplanted mice. n= 5.^a,b^ **I.** Representative bone histomorphometry (H&E, Tartrate-resistant acid phosphatase (TRAP), and Alkaline phosphatase (ALP) staining) images of the femurs with quantification of relative intensity of TRAP-positive osteoclasts and ALP-positive osteoblasts. n= 5 for TRAP staining, 3 for ALP staining. Scale bar, 800 µm (H&E staining) or 100 µm (TRAP and ALP staining). ^a,b^ ^a^Data shown are means ± SEM. The data presented in D and F represent the combined data from at least two independent datasets. ^b^*p < 0.05; **p < 0.01; ***p < 0.001; ****p < 0.0001 by one-way analysis of variance or two-tailed Student’s t testing.

### Increase of circulating eNAMPT through hypothalamic-adipose inter-tissue communication underlies the systemic aging retardation in the htNSC-grafted mice

To attain mechanistic hints behind the improvement in age-associated behaviors in the hypothalamus-transplanted mice, we first measured biomolecules associated with body metabolism/aging in circulating blood. Among those tested using a cytokine array kit, leptin levels were lower in the grafted mice compared to the control mice (Suppl. Fig. S8A–B), which was further validated by ELISA-based quantification (Suppl. Fig. S8C). This finding aligns with the established positive correlation between plasma leptin levels and body fat mass ^51,52^. The reduced levels of leptin also correspond to the increased bone density observed in the grafted mice ^53^. Lipid blood chemistry analysis showed dramatic reduction of LDL-C but no changes in the other factors examined (Suppl. Fig. S8D-G).

Additionally, we observed significantly higher levels of extracellular nicotinamide phosphoribosyltransferase (eNAMPT), an anti-aging factor secreted from adipose tissues ^54–56^, in the circulating blood of the grafted mice (Fig. 5A). The mRNA expression of NAMPT was also significantly elevated in the iWAT and eWAT of the hNSC-grafted mice (Fig. 5B). Treatment with a NAMPT inhibitor (FK-866, administered intraperitoneally, dosage: 10 mg/kg/day) or beta-adrenergic (SNS) blocker (propranolol, administered via drinking water, dosage: 0.5 g/L) two weeks post-transplantation (Fig. 5C), largely negated the anti-aging effects of the hNSC transplantation, both in terms of aging-related physiological declines (Fig. 5D-J) and histologic changes (Fig. 5K), indicating that the increased eNAMPT and/or symphathetic tone in peripheral tissue and blood plays a crucial role in the anti-aging effects observed in the htNSC-transplanted mice.

**Fig. 5.**
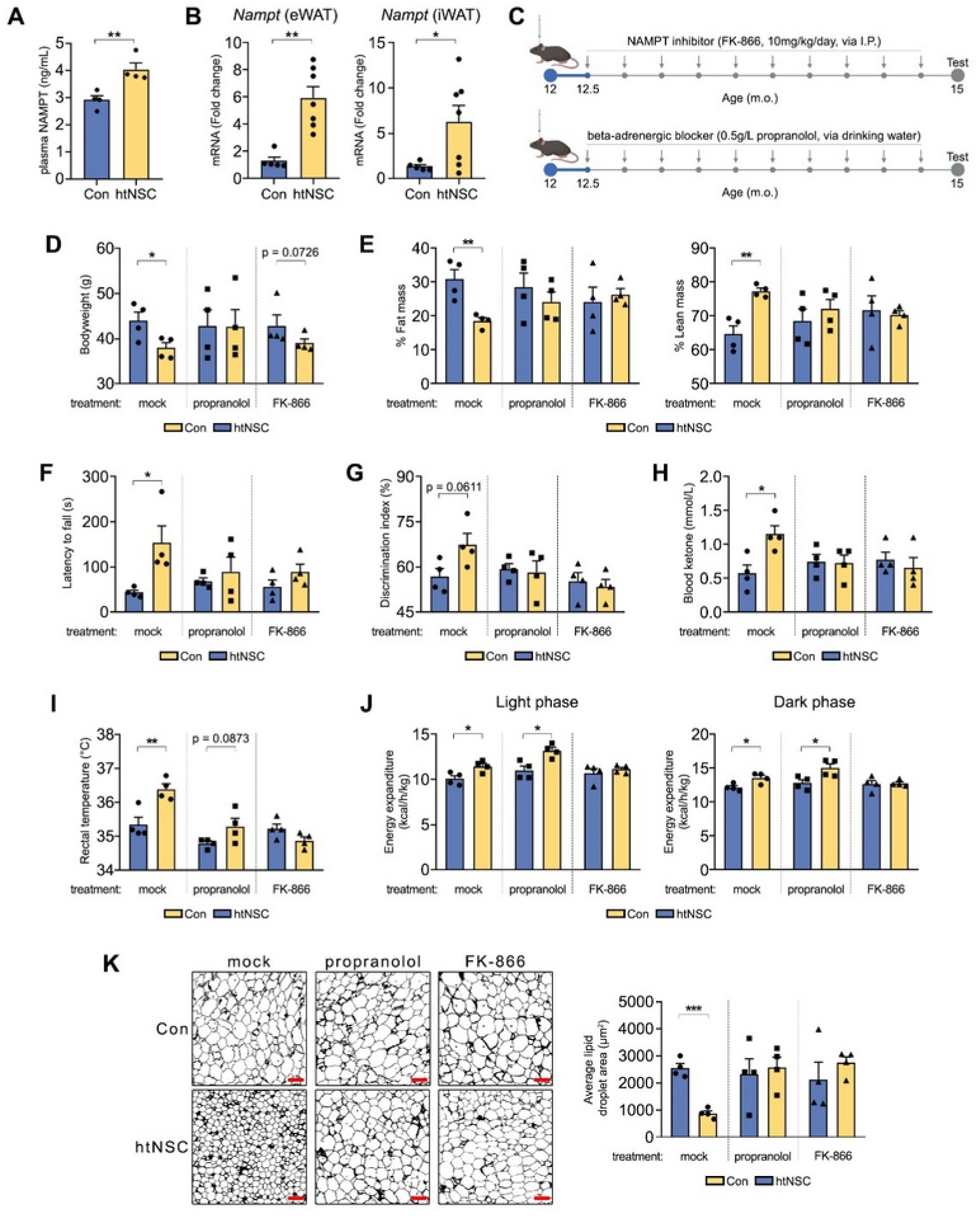
NAMPT and beta-adrenergic inhibitors reversed the anti-aging effects of the hNSC transplantation. **A.** ELISA-based determination of serum NAMPT levels in the transplanted and control mice. n=4. ^a,b^ **B.** mRNA expression of *Nampt* in iWAT and eWAT tissues assessed by qRT-PCR. n= 5 (control), 7 (transplanted). ^a,b^ **C.** Experimental scheme for inhibitor treatment in control and transplanted mice. Treatment began two weeks post-transplantation and continued for 2.5 months. NAMPT inhibitor (FK-866, 10 mg/kg) was administered intraperitoneally once a day. Beta-adrenergic blocker (propranolol, 0.5 g/L) was administered via drinking water. All analyses were conducted after 2.5 months inhibitor treatment. **D–E.** Bodyweight measurements (D) and NMR-based measurement of the fat and lean masses. n= 4. ^a,b^ **F-G.** Muscle strength and cognitive function assessment by grip test and Y-maze test, respectively. n= 4. ^a,b^ **H.** Fasting blood ketone levels. n= 4. ^a,b^ **I.** Rectal temperature. n= 4. ^a,b^ **J.** Energy expenditure measured in metabolic cages. n= 4. ^a,b^ **K.** Quantified lipid droplet areas (right) with representative H&E-stained tissue section images (left) of the iWAT. n= 4. ^a,b^ ^a^Data shown are means ± SEM. ^b^*p < 0.05; **p < 0.01; ***p < 0.001; ****p < 0.0001 by one-way analysis of variance or two-tailed Student’s t testing.

Furthermore, our blood examination revealed that circulating levels of ketone bodies were significantly elevated in the transplanted mice (Fig. 5H) and reversed upon NAMPT inhibitor treatment or beta-adrenergic blocker, suggesting that the increase in ketone bodies may also mediates the anti-aging. Conversely, this rise in ketone bodies may act as a factor to correct functional deficits and inflammation in the aged hypothalamus ^57,58^, indicating a potential positive feedback loop between elevated ketone body levels and improved hypothalamic function and inflammation. Ketone bodies, which increase in the blood during calorie restriction, are recognized as anti-aging metabolites ^59–61^. These findings collectively indicate that the htNSC-grafted hypothalamus stimulates adipose tissue to synthesize and release eNAMPT into systemic circulation via the sympathetic nervous system, thereby retarding systemic aging.

### Single-cell transcriptome analysis of aged hypothalamic brains grafted with htNSCs

To identify the transcriptional profile of the grafted hypothalami, we performed a droplet-based single-nucleus RNA-seq (snRNA-seq) analysis of cell nuclei extracted from the transplanted hypothalamus and control groups 4 months post-transplantation. We sampled six hypothalami (three per group) on four independent sequencing runs (two runs per group) (Fig. 6A). After filtering and doublet removal, 12,161 nuclei were clustered and projected onto a two-dimensional UMAP space for visualization (Fig. 6B). We identified 30 clusters: 23 neuron (10 clusters of GABAergic neurons and 13 clusters of glutamatergic neurons), two tanycyte, one astrocyte, one oligodendrocyte, one oligodendrocyte precursor cell (OPC), one microglia, and one pericyte cluster. For the sake of simplicity and general visualization, the 23 neuronal clusters were merged into glutamatergic and GABAergic neuron clusters, as described above, and the two tanycyte clusters were merged into one (Fig. 6B and Suppl. Fig. S9A). The cells were annotated manually based on well-established cellular markers: *Snap25* for pan-neurons, *Gad1* for GABAergic neurons, *Slc17a6* for glutamatergic neurons, *Agt* and *Slc1a3* for astrocytes, *Mbp* and *Plp1* for oligodendrocytes, *Olig2* and *Pdgfra* for OPCs, *P2ry12* and *Cx3cr1* for microglia, *Rax* and *Col23a1* for tanycytes, and *Slc4a7* and *Flt1* for pericytes (Fig. 6C and Suppl. Fig. S9B). Many well-known hypothalamus-specific neuronal subtypes were identified in the neuron subclusters. For example, GABA neuron subcluster 2 (GABA-2), which contained galanin-secreting neurons, are highly enriched in the hypothalamus and could be involved in feeding behavior ^62,63^, and Glut-11 is a melanin-concentrating hormone (MCH; gene name, *Pmch*) neuron, which is a sub-population of neurons in the lateral hypothalamus (Fig. 6D–E and Suppl. Fig. S10A). A gene ontology (GO) analysis showed that the top signature genes in each cluster were enriched in the GOs consistent with their known biological functions. For instance, GOs regarding ‘synaptic transmission’ and ‘neurotransmitter’ were specifically enriched in the neuronal clusters, and GOs regarding ‘immune process’ were specifically enriched in the microglial cluster (Fig. 6F). No significant difference in the number of cells in each cluster was found between the grafted and control groups (Fig. 6G).

**Fig. 6.**
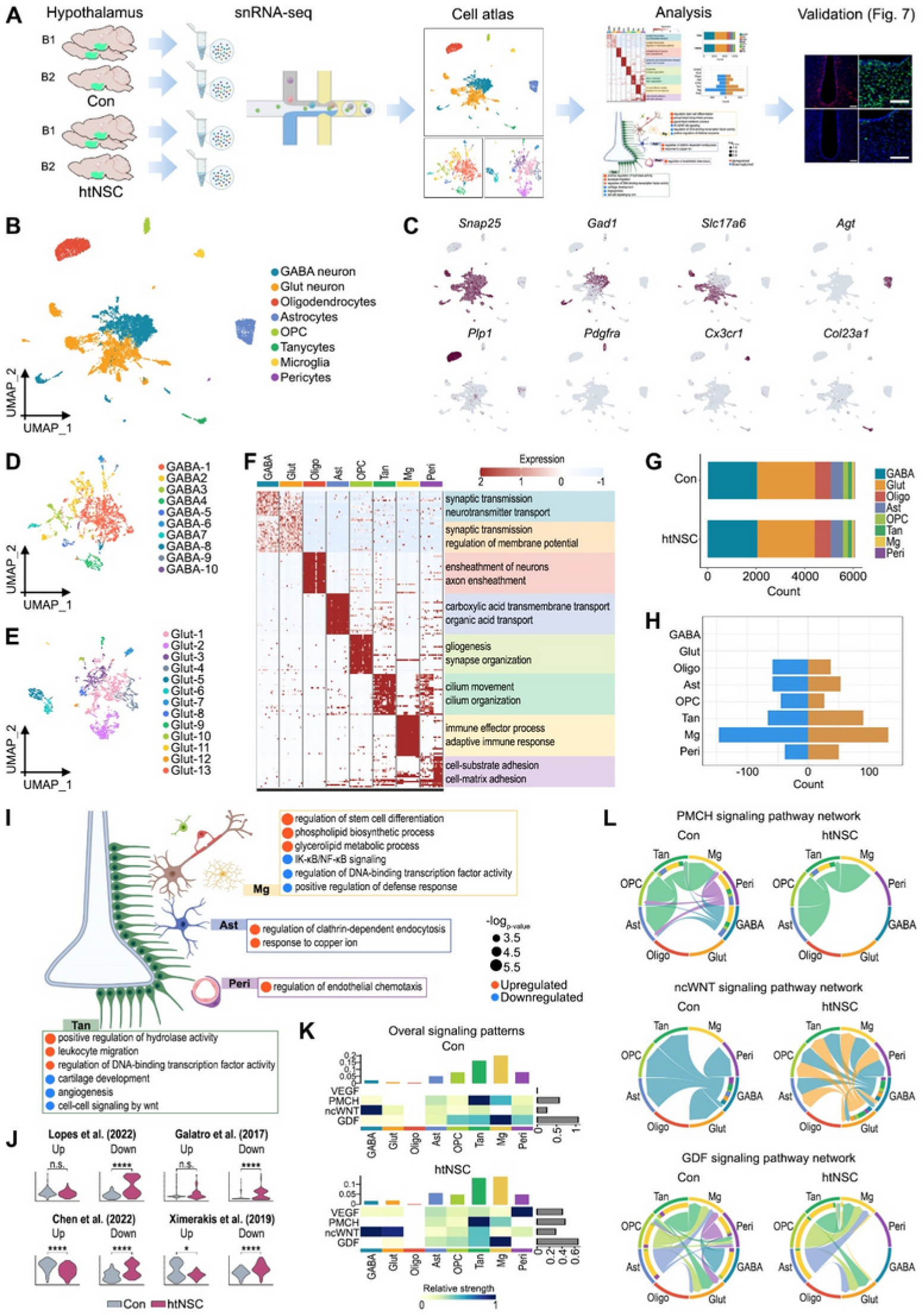
Single nucleus RNA-seq analysis of transplanted and control hypothalami. **A.** Schematic of the snRNA-seq experiment. **B–C.** UMAP plot of 12,161 nuclei filtered and processed from control and transplanted hypothalami (B) and feature plot showing representative cluster transcripts (C). **D–E.** UMAP plots of the GABA-neuron subcluster (D) and Glut-neuron subcluster (E). **F.** Heatmap showing the top 25 signature genes among the clusters and the two most representative GO results. **G.** Stacked bar plot showing the composition of each cell type in the dataset for each group. **H.** Bar plot showing the number of upregulated (orange) and downregulated (blue) transcripts in all cell types for the transplanted group. **I.** GO (biological process) enrichment analysis of DEGs from the non-neuronal clusters. **J.** Violin plot of module scores for microglial nuclei in gene sets obtained from various studies. The statistical analysis used the Mann-Whitney test. **K.** Subset of a heatmap of the cell-to-cell interaction network showing the overall (ingoing and outgoing) signal strength of VEGF, PMCH, ncWNT, and GDF signaling. **L.** Circle plots of an inferred signaling network of PMCH, ncWNT, and GDF signaling.

Next, we performed a differential gene expression (DEG) analysis on each cluster to examine the cell-specific changes that could be mediating the anti-aging effect of the intra-hypothalamic htNSC-transplantation. Interestingly, no statistically significant DEGs were detected in the pan-GABA and pan-Glut clusters (Fig. 6H). By contrast, abundant DEGs were detected in non-neuronal cell clusters. Among them, the microglia showed the most dramatic gene expression changes following intra-hypothalamic htNSC transplantation, which was followed by tanycytes, astrocytes, OPCs, and oligodendrocytes. These findings are consistent with the previous report that glia-specific genes predict age with greater precision than neuron-specific genes ^64–66^.

The microglial DEGs that were downregulated in the grafted hypothalami were enriched in the GOs associated with immune response, specifically ‘IκB kinase/NF-κB signaling’ (Fig. 6I). Indeed, the expression of central microglial NF-κB subunits (*Nfkb1*, *Rel*, *Relb*, and *Ikbkg*) were dramatically downregulated in the htNSC-grafted group (Fig. 7E). This is consistent with the hypothalamic microinflammation theory of aging, which is in turn supported by a recent single-cell transcriptomic result about the aged hypothalamic median eminence ^67,68^. Specifically, it has been shown that hypothalamic Iκκb and NF-κB are molecules responsible for aging ^10,69^ and that htNSCs dampen NF-κB signaling and hypothalamic micro-inflammation in a paracrine manner ^70^. The microglial genes upregulated by htNSC transplantation were annotated to GOs related to ‘metabolic process,’ specifically ‘lipid metabolism’ (Fig. 6I). Consistently, it has been reported that dysregulated lipid metabolism and lipid accumulation in microglia represent a dysfunctional and pro-inflammatory state in the aging brain ^71^. We further compared gene expression changes in the microglia of grafted hypothalami (vs. vehicle-injected controls) with those of age-associated microglial genes recorded in multiple published datasets ^67,72,73^. Single-cell expression levels of the microglial genes downregulated during aging in all the published datasets (calculated using ‘AddModuleScore’ in the Seurat package) were significantly upregulated in the microglia of grafted hypothalami, compared with the levels in the vehicle-treated control hypothalami (p-value <0.05, statistical method Wilcoxon rank-sum test) (Fig. 6J). Concordantly, microglial genes with age-dependent upregulation in the datasets of Chen (2022) and Ximarekis (2019) were significantly downregulated in the microglia of the grafted hypothalami, together indicating that age-dependent microglial signatures were corrected in aged hypothalami by htNSC transplantation.

**Fig. 7.**
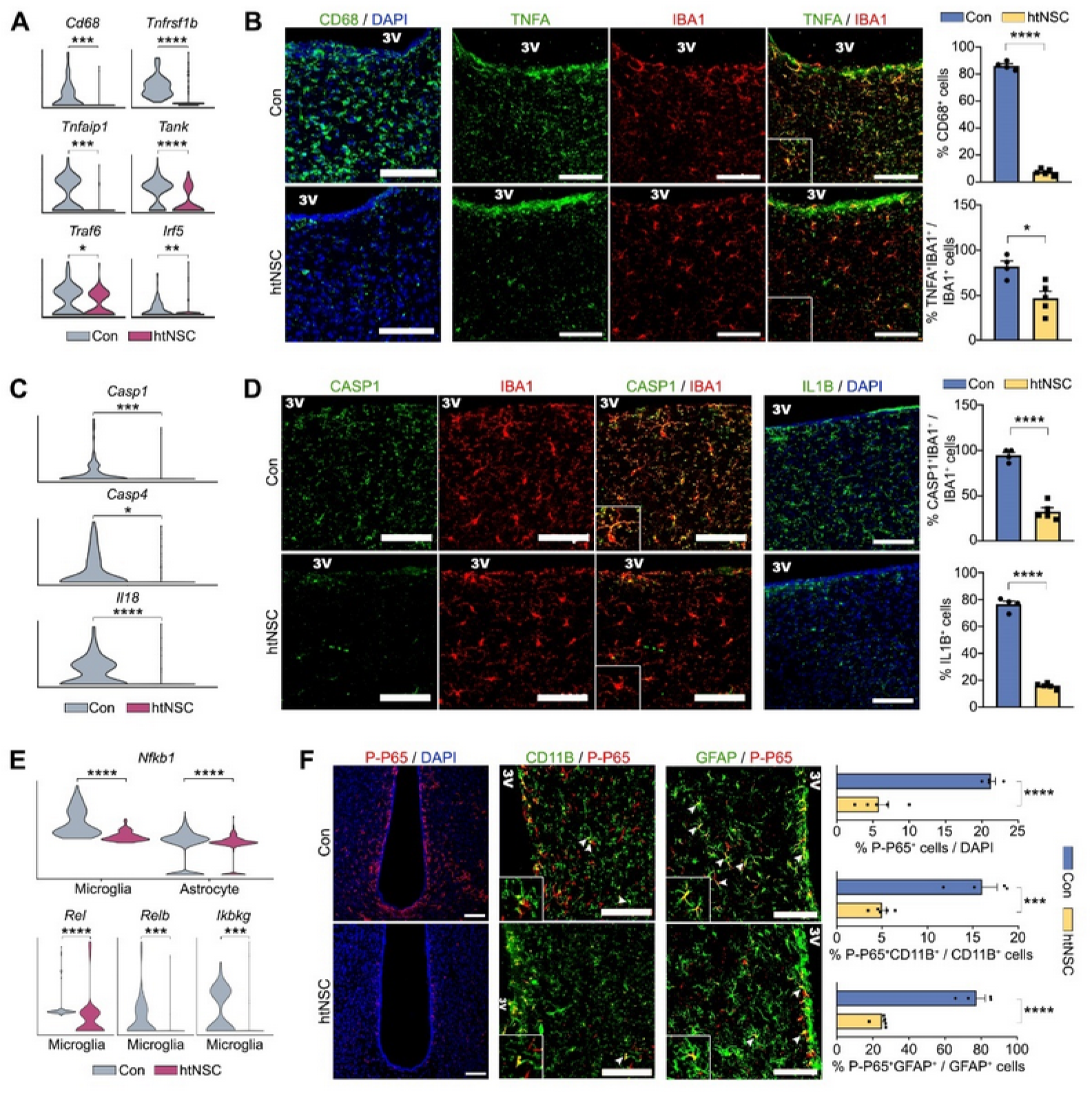
Neuro-inflammation in aged hypothalami subsided following htNSC transplantation. **A–B.** Hypothalamic inflammation, as estimated by the snRNA-seq analysis of inflammatory gene expression (*Cd68*, *Tnfrsf1b*, *Tnfaip1, Tank, Traf6,* and *Irf5*) in the microglial clusters (C) and the immunohistochemical analysis for the percentage of CD68+/DAPI+ and TNFA+/IBA1+ cells (D). Scale bars, 100 µm. n = 3 (control), 3 (transplanted) in (C)^c,d^ and n = 4 (control), 5 (transplanted) in (D)^a,b^. **C–D.** SnRNA-seq- and immunohistochemistry-based determination of NLRP3-inflammasome activation in the aged control and grafted hypothalami. Scale bars, 100 µm. n = 3 (control), 3 (transplanted) in (E)^c,d^ and n = 4 (control), 5 (transplanted) in (F)^a,b^. **E–F.** NF-κB signal activation in the aged control and grafted hypothalami, as assessed by snRNA-seq (A) and immunohistochemical (B) analyses. Violin plot of *Nfkb1* in microglial and astrocyte clusters and *Rel, Relb,* and *Ikbkg* in microglia from the snRNA-seq data (A). The percentage of p-P65+ in the total (DAPI+), microglial (IBA1+), and astrocytic (GFAP+) cell populations (B). Scale bars, 100 µm. n = 3 (control), 3 (transplanted) in (A)^c,d^ and n = 4 (control), 5 (transplanted) in (B)^a,b^. ^a^Data shown are means ± SEM. ^b^*p < 0.05; ***p < 0.001; ****p < 0.0001 by two-tailed Student’s t testing. ^c^Data shown are medians. ^d*^p < 0.05; **p < 0.01; ***p < 0.001; ****p < 0.0001 by Mann-Whitney U testing.

The tanycyte cluster had enrichment in GOs related to cell-to-cell and intracellular signaling (Fig. 6I). That finding is consistent with the role of naïve tanycytes as the portal between the hypothalamus and the rest of the body, the place where hormones and metabolic cues are exchanged. The most enriched GO in the tanycytes of the grafted hypothalami was ‘positive regulation of hydrolase activity.’ It has been demonstrated that after being released from hypothalamic neurons, pro-TRH requires hydrolase-mediated activation in tanycytes ^22,74^. Therefore, it is possible that htNSC transplantation also restores the tanycyte-mediated TRH signaling process that usually decreases during aging ^75^.

We further analyzed the htNSC transplantation–mediated DEGs in 23 neuron subclusters characterized by their unique gene expression signatures (Suppl. Fig. S10A). GABA-4 and GABA-6 were the GABAergic neuron clusters whose gene expressions were most highly modulated by htNSC transplantation, whereas Glut-5, Glut-6, Glut-7, and Glut-8 were the most highly affected among the glutamatergic neuronal clusters (Suppl. Fig. S10B). Among those, it is noted that GABA-4 was the *Vipr2*+ neuronal population associated with circadian clock, while GABA-6 was identified as *Lef1*+ neuron whose function is unknown ^76^.

A recent study concluded that Ppp1r17 neuron population in the DMH regulates the aging process in mice through activation of eNAMPT in the adipose tissues ^56^. Interestingly, we identified Glut-7 cluster as *Ppp1r17+* neuron which gene expression are highly modulated in the transplantation group (Suppl. Fig. S10A–B). We also observed significant increase in the c-FOS immunoreactivity in the DMH of the transplanted mice (Suppl. Fig. S10C) that is consistent with previous finding and explained the increased plasma eNAMPT ^56^.

Loss of cell-to-cell communication is one of the hallmarks of aging. Using known ligand– receptor interactions, it is possible to infer information flow from one cell type to another. A cell-to-cell communication analysis using CellChat ^77^ showed that pericyte VEGF signaling was restored by htNSC transplantation (Fig. 6K). Downregulation of the VEGF signal is a major underlying cause of age-dependent blood brain barrier (BBB) dysfunction ^78^. MCH is an orexic neurohormone that is mainly secreted by GABAergic neurons in the hypothalamus ^79^. Consistent with the increased body energy metabolism that we saw in the grafted mice, GABAergic neuronal output of MCH (*Pmch*) and neuronal communication with glial cells were diminished in the grafted hypothalami (Fig. 6L). Additionally, alterations in non-canonical WNT and GDF signal–mediated cell–cell communications were also manifested following htNSC transplantation.

### Neuro-inflammation in aged hypothalami corrected by intra-hypothalamic htNSC implantation

Next, we validated the snRNA-seq data indicating that hypothalamic inflammation subsided following htNSC transplantation by examining protein levels with immunofluorescence staining. Consistent with the snRNA-seq data, the abundance of IBA1+ microglia and GFAP+ astrocytes in the hypothalamus were not altered by htNSC transplantation (Suppl. Fig. S11A–B). No morphologic differences in microglia or astrocytes were found between groups (Suppl. Fig. S11A–B). However, a marked reduction of Cd68+ cells, microglial TNF-α, Casp1, and Il-1β expression in the treatment group were confirmed both in the snRNA-seq and in immunohistochemical analysis (Fig. 7A–D). More importantly, the number of phosphorylated p65 (activated p65) positive cells was also greatly reduced in the transplanted group, as well as its co-expression in microglia and astrocyte (Fig. 7E–F). Interestingly, the reduction of hypothalamic p65 was completely reversed by NAMPT inhibitor (Suppl. Fig. S12). Chronic hypothalamus inflammation has been identified as both cause and consequence of systemic metabolic dysregulation ^80–82^. Unexpectedly, propanolol treatment dramatically reduce p65 expression in both control and transplantation group (Suppl. Fig. S12), possibly through direct inhibition of microglial β-AR by propranolol ^83–85^.

## DISCUSSION

In this study, we demonstrated that htNSCs derived from hPSCs exhibit health-span-promoting benefits when transplanted into aged hypothalamus. The correction of age-associated changes was represented by increased energy metabolism and glucose tolerance, with upregulated browning/thermogenic/catabolic, and sympathetic adrenergic gene and protein expression. In addition to changes in their body/energy metabolism and physiology, the mice grafted with human htNSCs in this study and mouse htNSCs in the study of Zhang et al. ^10^ showed improved memory and cognitive functioning. Considering that the hypothalamus is the master coordinator of a myriad of homeostatic functions that are deregulated during aging, the anti-aging effects observed in this study are likely to reflect the central action of the grafted human htNSCs (or their differentiated progenies) in rejuvenating host hypothalamus cells or environment to allow systemic body metabolism and brain function to be more efficiently regulated. We identified hypothalamus-SNS-eNAMPT axis that is required for the anti-aging effect.

Mechanistically, intra-hypothalamic htNSC transplantation increased neuronal activity in the DMH, which activates the SNS to induce the synthesis and release of the anti-aging factor eNAMPT from adipose tissues. The elevated levels of eNAMPT in systemic circulation are believed to play a crucial role in enhancing body and lipid metabolism in the grafted mice ^56^. In addition, eNAMPT can be transported to the hypothalamus, where it promotes rejuvenation and reduces inflammation through NAD+ production and SIRT1 activation ^54,55^. Conversely, products such as lipids and pro-inflammatory cytokines, increased by dysregulated peripheral energy metabolism, cause chronic inflammation in the hypothalamus. ^86,87^. Consequently, the physiologic effects of htNSC-transplantation were abolished upon SNS or eNAMPT inhibition. These findings support the concept of ‘Inter-organ communication in aging’ and highlight a feed-forward mechanism between improved systemic metabolism and hypothalamic function in transplanted mice ^88^. In conclusion, this study has shown that implanting human htNSCs derived from hPSC-organoids holds therapeutic promise for promoting healthy metabolic network which may also slow aging process.

Although this study shed some insight on the potential of htNSC replacement therapy, this study has several limitations. In this study we only utilized male mice. Sexual dimorphism of aging is a known phenomenon that we could not cover in this study. Secondly, we also used only single strain of mice; therefore, it is unclear although highly probable that the result observed here can be extrapolated into another genetic background or other species.

## MATERIAL AND METHODS

### Reagents and resources

Detailed information of reagents and resources used in this paper can be found in Supplemental Table S1.

### Cell culture

hESCs (H9) and hiPSCs (CMC-hIPSC-011, CS-001) were cultured on Matrigel-coated plates in feeder-free conditions in mTeSR-1 medium (Stemcell Technologies, Vancouver, Canada). As schematized in Fig. 1A, human htNSCs were prepared by adopting the organoid-based NSC preparation method that was recently published by our group ^28^. First, to generate human hypothalamus-like organoids, hESCs or hiPSCs were seeded at a density of 5,000 cells/well in a low-attachment, 96-well, round-bottom plate in neural induction medium (N2:Neurobasal, 1:1, Thermo Fisher Scientific, Waltham, Massachusetts) containing 2% B27 without vitamin A (Thermo Fisher Scientific), 200 µM ascorbic acid (MilliporeSigma, Burlington, Massachusetts), and 25 mg/L insulin, (Thermo Fisher Scientific). The ROCK inhibitor Y27632 (20 μM, MilliporeSigma) was added for 24 hours after cell seeding. Hypothalamus patterning of the differentiating hESCs/hiPSCs was induced by adding SB431542 (10 μM, Tocris, Bristol, UK) and noggin (200 ng/ml, Peprotech, Cranbury, New Jersey) for days 0–7 and adding sonic hedgehog (SHH, 100 ng/mL, Peprotech) and purmorphamine (2 μM, MilliporeSigma) for days 1–13. The medium was replenished every other day. Floating spheres (organoids) were transferred into low attachment, 6-well plates (16 spheres/well) on day 13 in neural expansion medium (N2 containing 16 µg/L insulin, 200 µM ascorbic acid, 100 U/ml penicillin G, and 100 μg/ml streptomycin) supplemented with bFGF (20 ng/mL, Peprotech), a mitogen for NSC, and cultured on an orbital shaker (80 rpm) until day 20. The organoids enriched with htNSCs were chopped, dissociated with Acutase (Stemcell Technologies), and plated on poly-L-ornithine/fibronectin-coated plates in bFGF-supplemented neural expansion medium. The htNSCs were expanded with cell passages at 5–7-day intervals. Y27632 was added for only 1 day after each cell passage. Terminal differentiation of the htNSCs was induced in N2 medium supplemented with 200 µM ascorbic acid, 10 ng/mL BDNF (Peprotech), 10 ng/mL GDNF (Peprotech), 1 μM db-cAMP (MilliporeSigma), and penicillin/streptomycin. As a control, NSCs with ventral midbrain identity (ventral midbrain-type NSCs; VM-NSCs) were prepared as described previously ^28^.

### Animal care and study approval

Middle-aged C57BL/6J male mice (11–12 months) were purchased from the Korean Basic Science Institute (Gwangju, South Korea) or Jackson Laboratory (Yokohama, Japan) and young-aged C57BL/6J male mice (2–3 months) were purchased from Korean Basic Science Institute (Gwangju, South Korea). All mice housed in a specific pathogen-free barrier facility with a temperature of 23 ± 1°C, a humidity of 50 ± 10%, a 12 h light/dark cycle, and free access to food (standard chow, 5053 PicoLab Rodent Diet 20, Lab Diet, Saint Louis, Missouri) and water. All procedures for the animal experiments were approved by the Hanyang College of Medicine Institutional Animal Care and Use Committee (IACUC) (approval number, 2018-0217A and 2020-0142A) and the Korean Institute of Science and Technology (KIST) IACUC and Institutional Biosafety Committee (Approval number, KIST-2019-048).

### Transplantation procedures

For cell transplantation, 12-month-old mice received stereotaxic surgery under anesthesia induced by a mixture of Zoletil (37.5 mg/kg) and Rompun (5.83 mg/kg) in isotonic saline. Human htNSC dissociated into single-cells using Accutase and subsequently suspended in 1μl of N2 medium (1 × 10^5^ cells/µL) were injected into the MBH (AP: –1.5 mm, ML: ±0.4 mm, DV: – 5.6 mm) at a rate of 0.25 µL/minute for 2 minutes using 26G needle, and the injector was then left in place for an additional 5 minutes before removal. A sham operation with an N2 medium injection was carried out for the control group. The animals received daily injections of cyclosporine A (10 mg/kg, i.p.) starting 1 day before the grafting and continuing for 3 weeks.

### Behavioral analyses

All behavioral tests were performed in the behavioral testing room. An ANY-maze video-tracking system (Stoelting, Wood Dale, Illinois) equipped with a digital camera connected to a computer was used. The following behavior tests were conducted in compliance with the ARRIVE guidelines. Detailed analyses in the Supplemental Methods.

### Measurement of systemic body metabolism

Assays for body composition, metabolic cage experiment, glucose tolerance test, body temperature, bone density measurement, and bone histomorphometry are described in the Supplemental Methods.

### Histological analysis

Mice were transcardially perfused with 30 mL of PBS (pH 7.4), followed by 30 mL of 4% paraformaldehyde (PFA) in PBS. Their brains were dissected out and post-fixed with 4% PFA overnight at 4°C and then transferred to 30% sucrose for 7 days for cryoprotection. The cryoprotected brains were frozen in TissueTek OCT compound (Sakura Finetek, Torrance, California) and sliced into 30 µm sections using a cryostat (Leica, Nussloch, Germany). For peripheral tissues, the WAT, BAT, livers, and femurs were dissected from the mice without fixation and divided into two halves for histochemical and Western blot analyses. The tissues for histochemical staining were fixed overnight at 4°C in 4% PFA in PBS, paraffin embedded, and sectioned into 4 μm slices that were stained with hematoxylin and eosin (H&E) following the standard procedure.

### Determination of protein and mRNA expressions

Protein expressions in the htNSC cultures and mouse hypothalamic tissues were assessed using immunostaining and Western blot analyses. Serum levels of cytokines and leptin were also determined using a Proteome Profiler Mouse XL Cytokine Array and a Mouse/Rat Leptin Quantikine ELISA Kit (R&D Systems, Minneapolis, Minnesota), respectively. mRNA expressions were assessed using quantitative PCR, global, and single cell transcriptomic analyses. Detailed methods are described in the Supplemental Methods.

### Statistical analysis

No statistical method was used to predetermine sample sizes, which were chosen with adequate statistical power based on literature and experience. Data shown from representative experiments were repeated with similar results in at least 3 independent experiments, unless otherwise indicated by sample size. Homogeneity of variance was tested using F-test for two groups, and Brown-Forsythe test for more than two groups. Data were analyzed using two-tailed unpaired Student’s t-testing, one-way analysis of variance (ANOVA), or Mann-Whitney U-testing. Welch’s correction was applied for datasets with unequal variance. Tukey post-hoc test was used for one-way ordinary ANOVA test while unpaired t-testing was used for one-way Welch ANOVA test. All multiple testing p-values were adjusted using Benjamini and Hochberg correction. For behavioral analyses, all data points below or above mean ± 1.645 * SD (corresponding to 5% in the normal distribution) are considered as outliers and were removed from statistical analyses. Additional analyses using linear mixed modeling and Monte-Carlo permutation (n = 10,000) were applied in behavior datasets. In all cases, results were considered statistically significant when p < 0.05. Error bars show the mean ± standard error of the mean (SEM) or the median, and p values are represented as: *p < 0.05, **p < 0.01, ***p < 0.001, and ****p < 0.0001. Statistical analyses were performed with Prism software version 8 (GraphPad Software) or R.

## Data availability

The datasets produced in this study are available in the following databases:

hPSCs-derived htNSC bulk RNA-seq: GSE214877

hPSCs-derived htNSC organoid scRNA-seq: GSE215174

htNSC-transplanted aged mouse hypothalamus snRNA-seq: GSE215174

Values for each data point presented in this paper can be found in the Supporting Data Values XLS file.

## Author contribution

S.H.L. and M.S.K. supervised the project. S.H.L., M.S.K., Y.A.S., Y.L., and K.P. designed experiments, and drafted and revised the manuscript. Y.A.S., Y.L., K.P., S.H., J.K., M.J.S., N-K.L., K-S.P., T.H., S.C., K.W.K., D.J.Y., W-Y.P., K.Y.H., S.G.Y., I.Y.K., and J.K.S. performed experiments. Y.A.S., Y.L., and K.P. analyzed and interpreted data, and performed statistical analyses. S.H.L., M.S.K., Y.A.S., J.K.S., T.Y.L., and M.S.K. acquired funding.

