## Supplemental File for "Human hypothalamic neural stem cells transplantation ameliorates age-associated dysfunction through SNS/eNAMPT axis activation in aged mice"

### Supplementary Figure

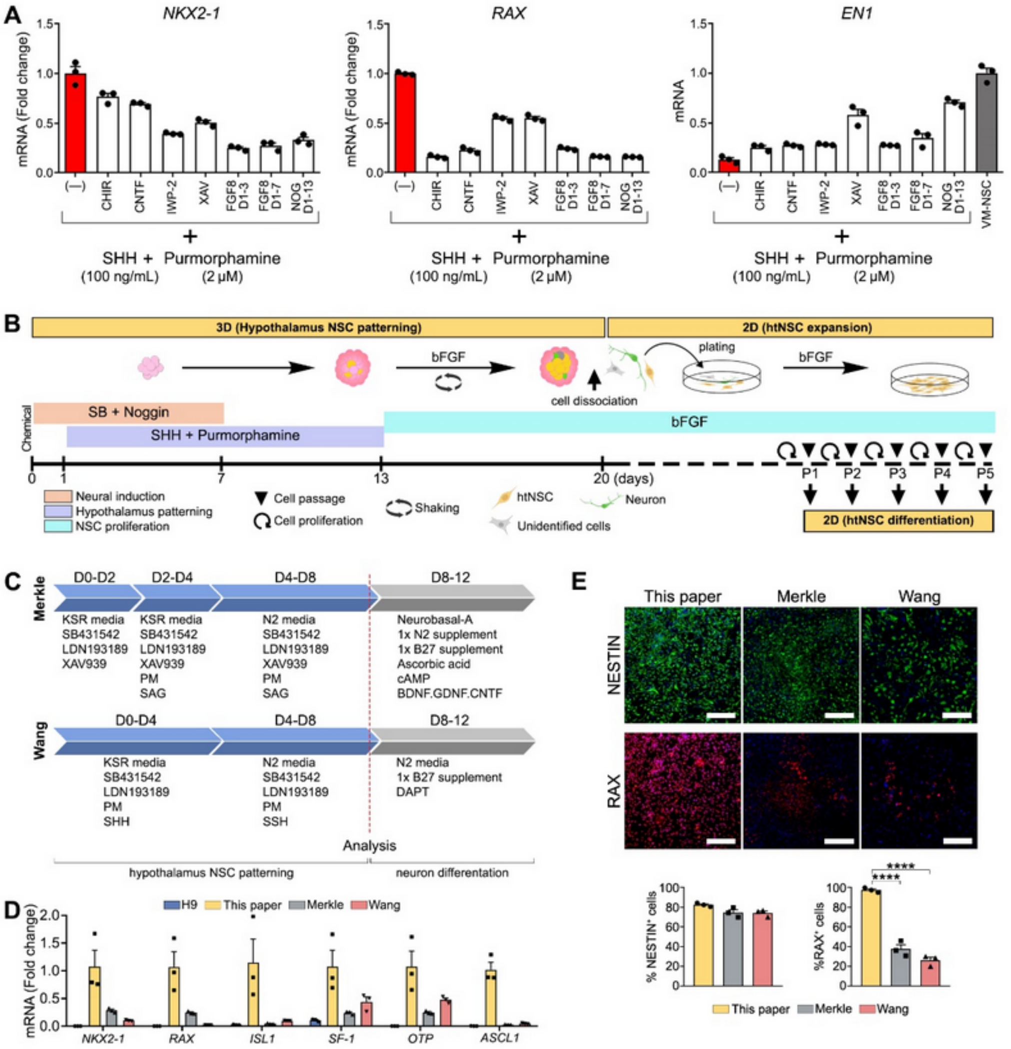


### Fig. S1. Optimization and validation of hypothalamus patterning in hPSC-derived htNSC cultures

**A.** The hypothalamic patterning in the differentiating hESC (H9) cultures treated with the chemicals indicated (plus SHH + purmorphamine) was judged as high expression of hypothalamus-specific markers (*NKX2-1* & *RAX*) and low expression of a midbrain-related marker (*EN1*) in a qPCR analysis. n=3. [a,b]

**B-E.** Comparison of hypothalamus-specific identity in the htNSC cultures derived by our optimized protocol (schematized in B) from other published protocols from Wang et. al. [1] and Merkle et al. [2] (schematized in C).

**D.** mRNA expression of the htNSC-specific marker genes (*NKX2-1*, *RAX*, *OTP*, *ISL1*, *SF1*) and neural progenitor cells marker (*ASCL1*) assessed by qRT-PCR. n=3. [a,b]

**E.** Immunocytochemistry-based quantification of NESTIN+ and RAX+ htNSCs. Scale bars, 100 µm. n=3.[a,b]

Data shown are means ± SEM. *$p < 0.05$; **$p < 0.01$ by one-way analysis of variance or two-tailed Student's t testing.

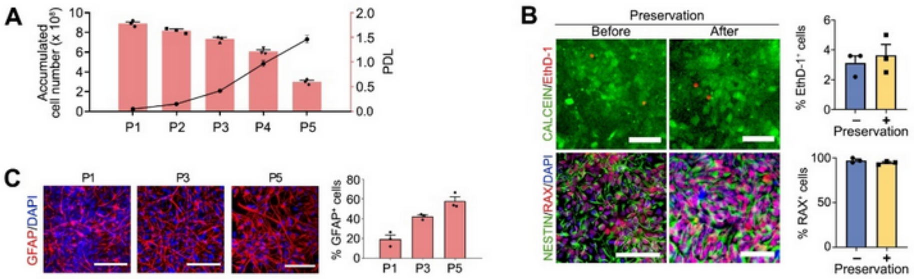


**Fig. S2 Maintenance of htNSCs identity and function during extended culture and cryopreservation.**

**A.** Cell expansion, estimated using a cell growth curve (line) and the population doubling level (PDL, bar). PDL=log(N/No)/log2, where N is the number of cells at the end of each passage, and No is the number of cells initially plated. n=3.[a]

**B.** Storage of htNSCs in liquid N2 without altering cell viability (Calcein/EthD-1) or htNSC-specific marker expression (NESTIN/RAX). Scale bars, 100 µm. n=3.[a,b]

**C.** Cell passage–dependent change in the htNSC differentiation propensity. The astrogenic differentiation potential of the htNSCs gradually increase during the cell passages. Scale bars, 100 µm. n=3.[a]

[a]Data shown are means ± SEM.

[b]*p < 0.05; **p < 0.01; ***p < 0.001; ****p < 0.0001 by two-tailed Student's t testing.

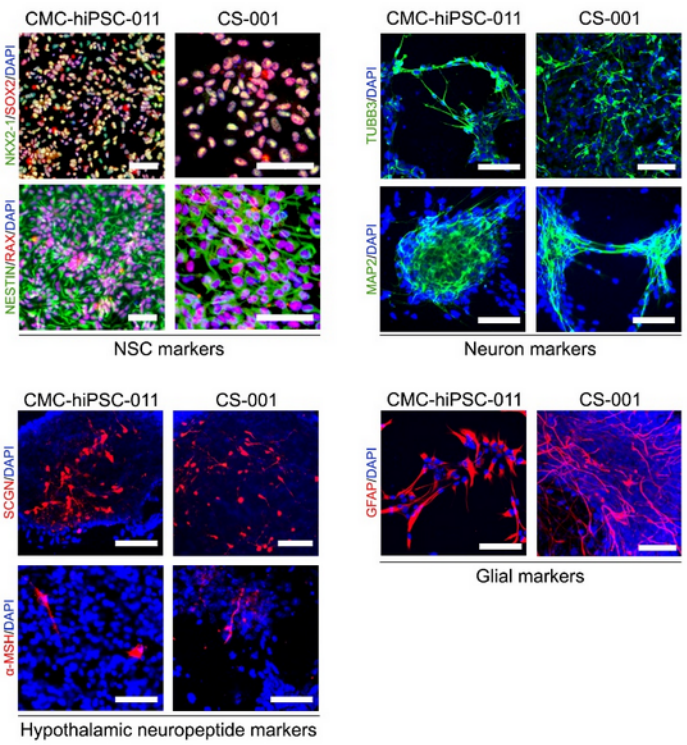


**Fig. S3. Derivation of htNSC cultures from the CMC-hIPSCs-011 and CS-001 hiPSC lines.** Representative immunostaining images for NESTIN+, RAX+, NKX2-1+, and SOX2+ NSCs at proliferation stage (expansion) and neurons expressing hypothalamus markers (TUBB3, MAP2, SCGN, and α-MSH) and astrocytes expressing GFAP 30 days after terminal differentiation. Scale bars, 100 μm.

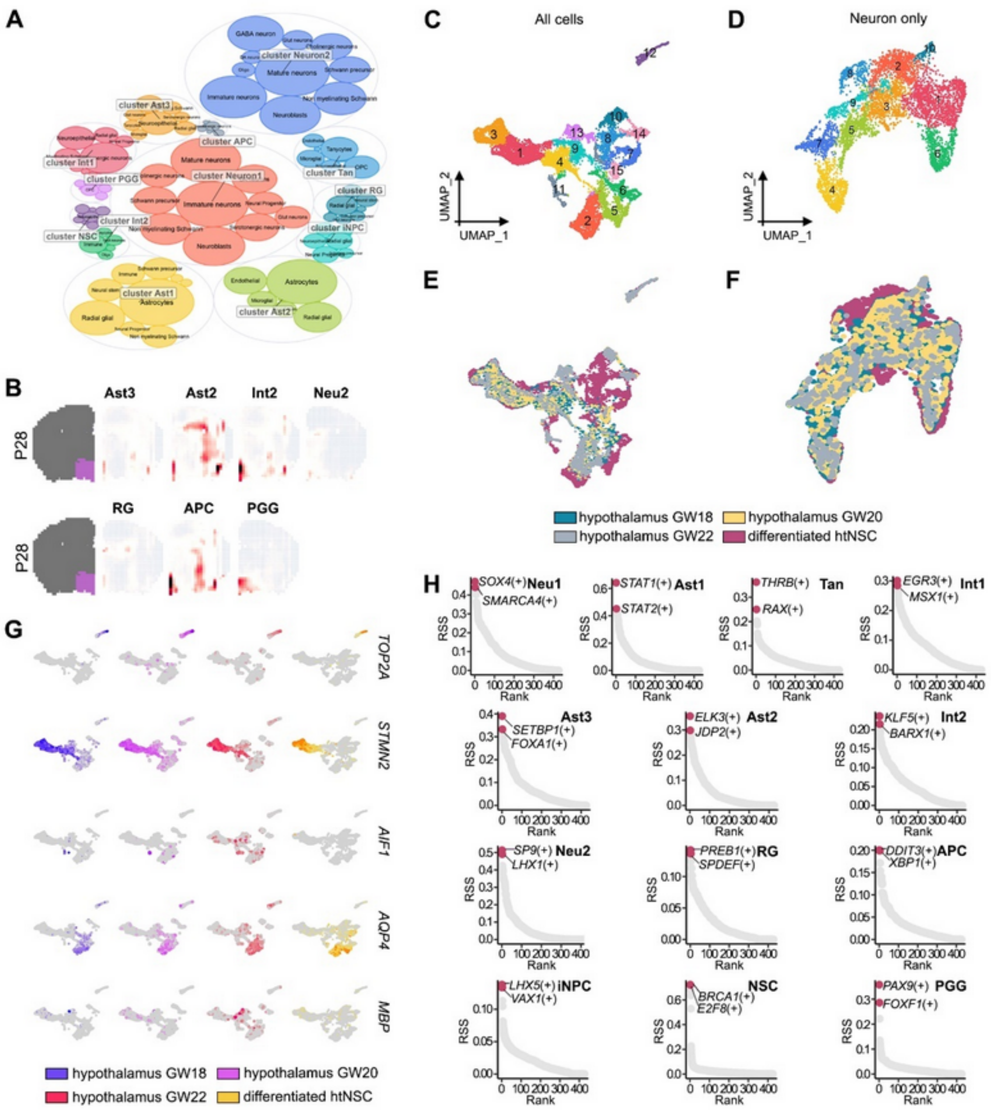


F

**Fig. S4. Validation of the cell clusters differentiated from the htNSC cultures.**

**A.** Automatic cluster identification using ScType.

**B.** Voxel-based spatial brain mapping of the clusters (Ast3, Ast2, Int2, Neu2, RG, APC, PGG) onto the mouse brain at postnatal day 28 (from the Allen Brain Institute).

**C-F**. UMAP plot of all cell clusters (C&E) and neuronal only clusters (D&F) from our differentiated htNSC cultures and published human fetal hypothalamus dataset (GW18-22) by cell types (C-D) and samples (E-F).

**G.** Features plots of representative cell-specific genes from the differentiated cell clusters.

**H.** Gene regulatory network analysis using pySCENIC to identify the potential upstream master regulators (regulons) in every cluster.

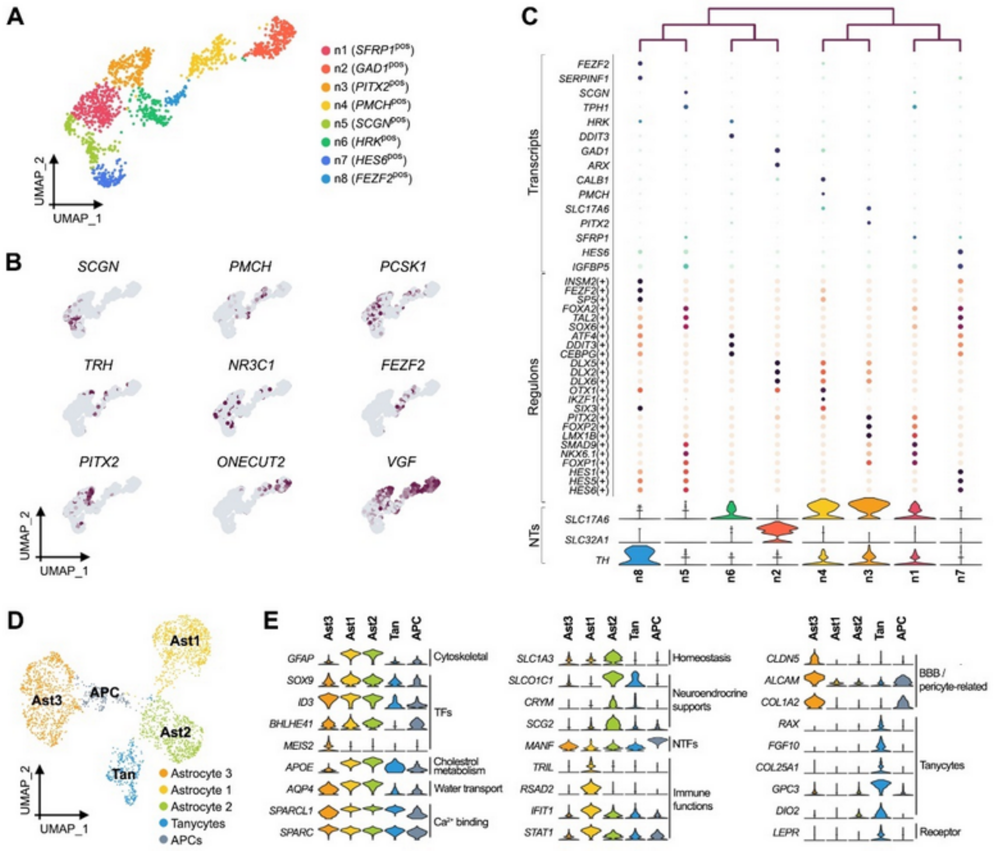


**Fig. S5. Re-clustering of neuronal clusters and astrocyte clusters.**

**A.** Re-clustering of the neuronal population into 8 subclusters of neurons.

**B.** Feature plot of representative hypothalamic neuropeptides (*SCGN*, *PMCH*, *PNOC*, *TRH*, *NR3C1*, *VGF*) and transcription factors (*FEZF2*, *PITX2*, *ONECUT2*).

**C.** Dot plots of selected transcripts (top) and regulons (middle) with violin plots of neurotransmitter phenotype gene expression (*SLC17A6*, *SLC32A1*, *TH*) (below) in each cluster of the neuronal subtypes.

**D–E.** UMAP plots of the astrocyte clusters (APC, Ast1, Ast2, Ast3, Tan) with the representative genes whose expression was enriched in each cluster.

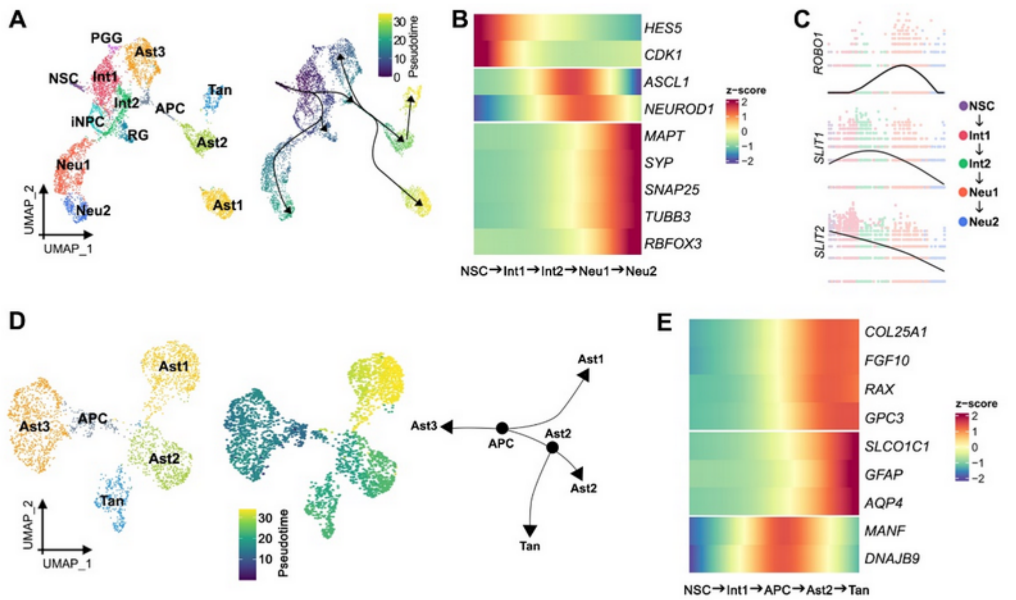


F

**Fig. S6. Pseudo-time trajectory analysis of the differentiated htNSC cultures.**

**A.** Monocle 3–based pseudo-time trajectory analysis of neuronal and glial cells differentiated from the hPSC-derived human htNSCs.

**B.** Heatmap of the cell-cycle genes (*HES5* & *CDK1*), early neuronal markers (*ASCL1* and *NEUROD1*), and late neuronal markers (*MAPT*, *SYP*, *SNAP25*, *TUBB3*, and *RBFOX3*) in the neuron trajectory (NSC, Int1, Int2, Neu1, Neu2) that were significantly associated with pseudo-time

**C.** Kinetic plot of *SLIT-ROBO* genes in the neuronal cell lineages (NSC, Int1, Int2, Neu1, Neu2).

**D.** Pseudo-time trajectory of the astroglial clusters (APC, Ast1, Ast2, Ast3, Tan).

**E.** Heatmap of astrocyte progenitor markers (*MANF* and *DNAJB9*), common astrocyte markers (*SLCO1C1*, *GFAP*, and *AQP4*), and tanycyte markers (*COL25A1*, *FGF10*, *RAX*, and *GPC3*) in the astroglial clusters that were significantly associated with pseudo-time.

A

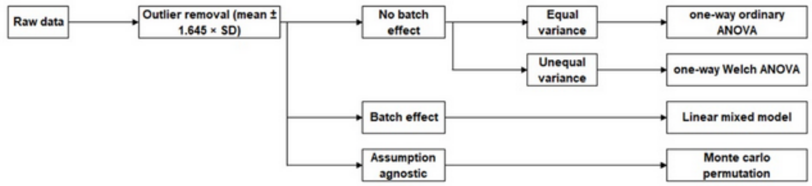


B

| | 13.5 month | | | | | | | | |
|---|---|---|---|---|---|---|---|---|---|
| | Young vs Con | | | Young vs htNSC | | | Con vs htNSC | | |
| BW | **** | **** | **** | **** | **** | **** | **** | **** | **** |
| GRIP | **** | *** | *** | ** | * | ** | *** | *** | *** |
| MWM | ** | ** | *** | ** | ** | *** | 0.8503 | 0.3417 | 0.8016 |
| NOR | 0.1818 | 0.1381 | * | 0.2972 | * | * | 0.5847 | 0.4662 | 0.7497 |
| OFT | 0.6136 | 0.4515 | 0.675 | 0.6136 | 0.0834 | 0.9583 | 0.6516 | 0.7546 | 0.7887 |
| ROTAROD | **** | **** | **** | *** | *** | **** | **** | **** | **** |
| TREADMILL | **** | **** | **** | *** | ** | **** | 0.1251 | * | 0.4269 |
| Y-MAZE | 0.4082 | 0.2288 | 0.56 | 0.4082 | 0.096 | 0.0552 | * | ** | 0.0552 |

| | 16 month | | | | | | | | |
|---|---|---|---|---|---|---|---|---|---|
| | Young vs Con | | | Young vs htNSC | | | Con vs htNSC | | |
| BW | **** | **** | **** | ** | *** | *** | **** | **** | *** |
| GRIP | ** | *** | ** | * | * | * | * | ** | ** |
| MWM | ** | *** | **** | 0.0562 | 0.2474 | ** | * | ** | * |
| NOR | **** | **** | **** | **** | *** | **** | * | *** | ** |
| OFT | 0.2972 | 0.3767 | 0.1897 | 0.747 | 0.1381 | 0.8803 | * | ** | * |
| ROTAROD | **** | **** | **** | 0.163 | * | ** | **** | **** | **** |
| TREADMILL | * | *** | **** | 0.8846 | 0.8357 | 0.3251 | ** | ** | ** |
| Y-MAZE | 0.9348 | 0.4577 | 0.7774 | 0.9348 | 0.7404 | 0.5965 | 0.9348 | 0.2665 | 0.9451 |

**Legend:** Linear Mixed Model
Monte Carlo
one-way Welch or ordinary ANOVA

F

**Fig. S7. Statistical analysis for the effect of intrahypothalamic htNSC transplantation on the physical and cognitive behaviors in aged mice.**

**A.** Diagram illustrating the process of data processing and statistical analyses applied to the behavior datasets.

**B.** Comparative table presenting statistical comparisons among three groups: control old mice (sham-operated), old mice with grafts, and young mice (aged 3-6 months), using three different statistical methods.

*$p < 0.05$; **$p < 0.01$; ***$p < 0.001$; ****$p < 0.0001$ by one-way analyses of variance, linear mixed model, or Monte Carlo permutation.

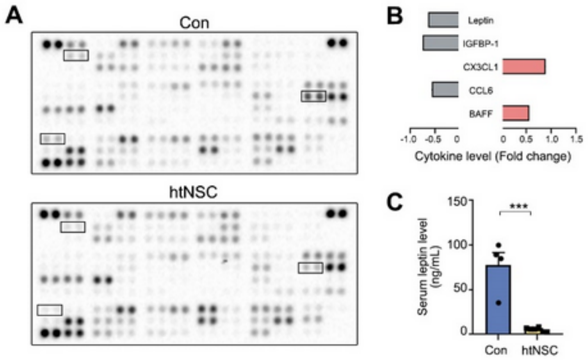


F

**Fig. S8. Serum analysis in the grafted (vs. control) mice.**

**A–B.** Array images of serum cytokine/chemokine profiles (A) and average cytokine/chemokine values with differences >1.5 fold (B) in the grafted and control mice. n= 2 (transplanted), 2 (control). Data shown are means.

**C.** ELISA-based determination of serum leptin levels in the transplanted and control mice. n= 6 (transplanted), 4 (control). Data shown are means ± SEM. ***p < 0.001; by two-tailed Student's t testing.

**D.**

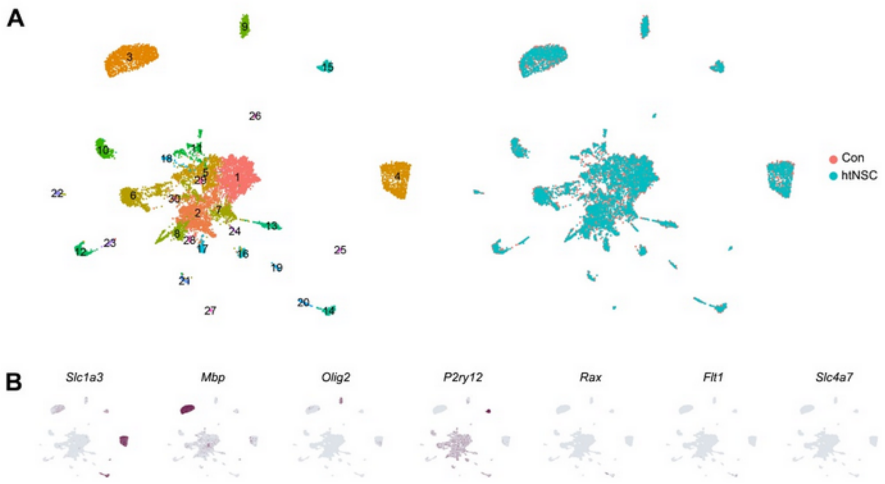


F

**Fig S9. Single nucleus RNA-seq analysis of transplanted vs. control hypothalami.**

**A.** UMAP plot of 12,161 nuclei from control and transplanted hypothalami projected into 30 clusters.

**B.** Feature plot showing additional representative cluster transcripts.

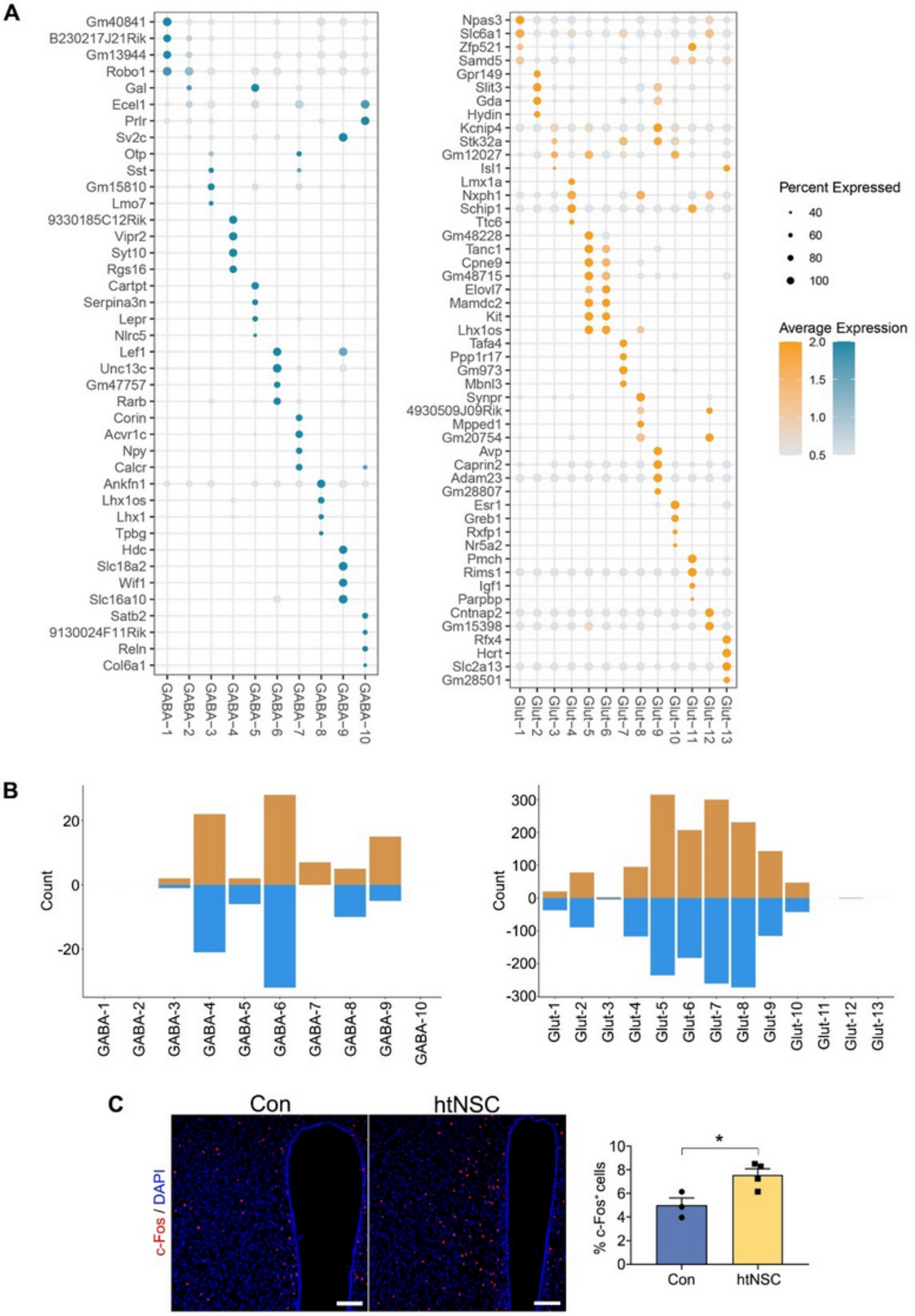


F

**Fig S10. Differential gene expression analysis of 23 neuronal subclusters.**

**A.** Dot plot showing the top 4 signature genes among the 23 neuronal subclusters.

**B.** Bar plot showing the number of upregulated (orange) and downregulated (blue) transcripts in the 23 neuronal subclusters of the transplanted group.

**C.** Neuronal activity estimated by the percentage of c-Fos-immunoreactive cells in the dorsomedial region of control and transplanted hypothalami. Data shown are means ± SEM. n = 3 (control), 4 (transplanted) mice. *$p < 0.05$ by two-tailed Student's t testing. Scale bars, 100 μm.

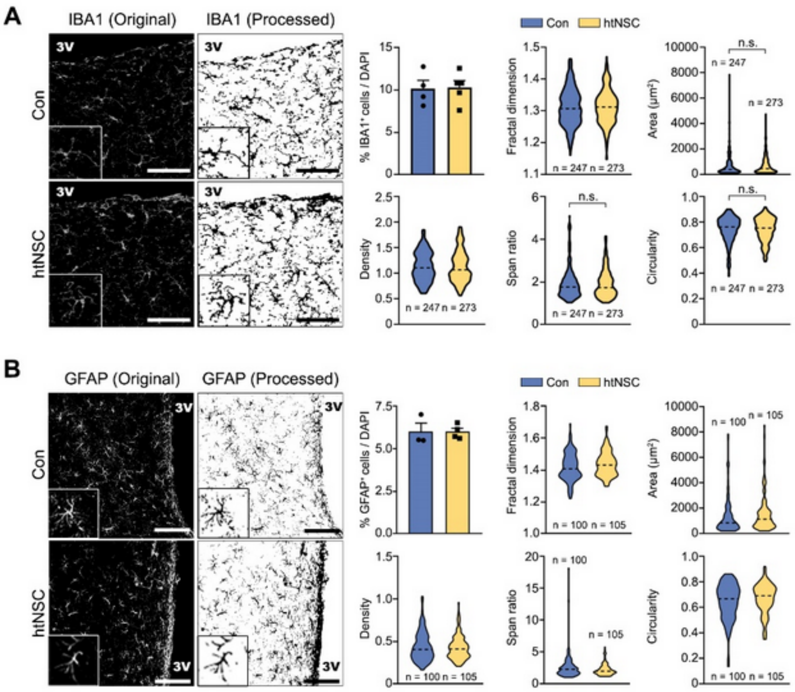


F

**Fig. S11. Glial cell numbers and morphologies in control and htNSC-grafted hypothalami.** The abundance of microglia (A, mice) and astrocytes (B, n = 3 (control), 4 (transplanted) mice) was assessed as the percentage of IBA1+/DAPI+ (A) and GFAP+/DAPI+ (B) cells. Data shown are means ± SEM. No significant difference was found in two-tailed Student's t testing. For the morphometric assay, fractal dimension, cell area, density (cell area/convex hull area), span ratio (ratio of major to minor axes of the convex hull), and cell circularity were measured in IBA+ (A, n = 247 (control), 273 (transplanted) cells) and GFAP+ (B, n = 100 (control), 105 (transplanted) cells) cell images processed in ImageJ. Data shown are medians. No significant difference was found in Mann-Whitney U testing. Scale bars, 100 μm.

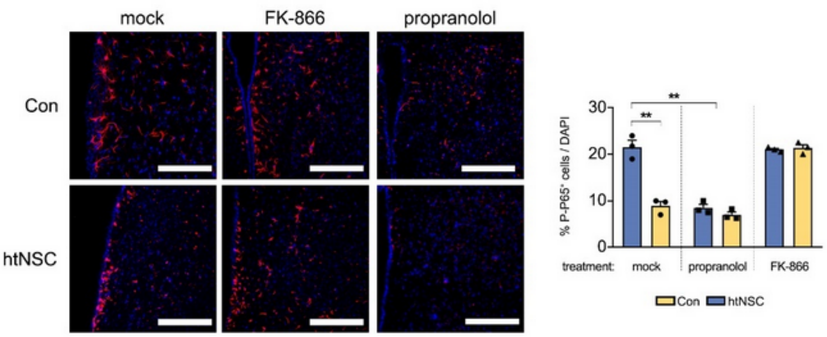


F

**Fig. S12. p-P65 immunostaining in mock, FK-866, and propranolol-treated control and htNSC-grafted hypothalami.**

Data shown are means ± SEM. n = 3. *$p < 0.05$ by two-tailed Student's t testing. Scale bars, 100 µm.

#### Supplementary method

##### Behavioral analyses

###### *Grip test*

The muscle endurance grip test was conducted as previously described [3]. Each mouse was lifted by the tail and placed on a homemade square grid (1-cm mesh size) that was then inverted 30 cm above a soft pad. The mouse was allowed to hang by his four limbs, depending on his age. The time that the mouse was able to hang was recorded. Three repeats were performed for each mouse with at least one hour rest between trials.

###### *Rotarod test*

The mice were trained on a rotarod (B.S. Technolab, Seongnam, South Korea) at a constant speed of 5 rpm for 60 s. After a 10-min rest, each mouse was placed on the rotarod at 5 rpm, and then the speed was accelerated by 5 rpm per min for 5 min. The time until the mouse fell was measured 3 times with an 1 hourrest period between trials.

###### *Treadmill*

The mice were warmed up to protect against running injury and failure before the experimental running. For acclimatization to a treadmill (Jeung-Do Bio & Plant, Seoul, South Korea), each mouse was placed on the treadmill at a low speed (5 rpm) for 5 min. The speed was then increased by 1 rpm every min for 25 min. The time at which the mouse retired was recorded.

###### *Novel object recognition*

The novel object recognition test was conducted to evaluate recognition-memory deficits. The mice were habituated to an open-field box (40 cm length × 40 cm width and × 50 cm height) for 10 min before the experimental sessions. After habituation, the mice were allowed to freely explore two identical objects in the open-field box for 20 min. Twenty-four hours after that, Mice was allowed 10 min for habituation and 10 min familization. After that, one of the two objects was replaced by a novel object, and mice were again allowed to freely explore the two objects for 10 min. The time spent exploring the familiar and novel objects was recorded. The preference index was calculated by dividing the time spent exploring the familiar or novel object by the total exploration time.

***Morris water maze***

The MWM test was used to measure spatial perception. The water tank was placed in the center of a small room with extra-maze cues (various shapes on a white background) and filled with 20-22°C water. Crayola non-toxic paint was added to make the water white and opaque. The diameter of the tank was 90 cm, and it was divided into four quadrants (northwest, northeast, southwest, and southeast). A circular platform with a diameter of 10 cm was placed 20 cm from the wall of the tank. Each mouse was placed in the water in the same starting location for all trials, and his latency and speed, to the platform were measured. For the hidden-platform training, the mice were first required to swim to a circular platform that was visible at 0.5 cm above the water level. The mouse had a 10 sec rest on the platfrom. If the mouse could not find the platform within 90 s, it was gently guided to the platform by hand. Each mouse received four consecutive days of training, consisting of one trial per entry location (opposite and left side quadrant of the platform quadrant) for a total of two trials per day. The platform was made invisible by submerging it 0.5 cm below the surface of the water. The mice were expected to find and rest on the invisible platform, and the latency to reach the platform were measured. On day 5, each mouse was subjected to a single probe trial, in which the platform was removed, and the animal was allowed to swim for 90 s. The amount of time spent in all quadrants, \ and swimming speed were measured. The learning and memory of the mice were indicated by the occupancy time (total time spent in the target quadrant) in their probe trials.

***Open-field test***

The open-field test was performed as previously described, with minor changes [4]. The mouse's movements were recorded for 10 minutes using a digital camera connected to ANY-maze animal tracking system software (Stoelting). The total distance traveled (meters) and time spent in the center/edge of the open field (seconds) were also recorded.

***Y-maze test***

The maze consisted of three arms, each 40 cm in length, forming 120-degree angle. Different shape cues were presented at the end of the arms. Mouse was placed at the end of one of the arms and freely explored for 8 minutes. An arm entry was considered when the mouse body completely enters the arm. It was counted the number of spontaneous alternations entering the new arm different from the previous arm. The percent of spontaneous alternation was calculated that the number of spontaneous alternations divided by the total number of entries minus 2 and multiplying by 100. (% spontaneous alternation = the number of spontaneous alternations / (the total number of entries-2) * 100) Mouse movement was tracked by ANY-maze tracking system (Stoelting).

**Body composition**

The total fat and total lean mass of the mice that received transplants were measured by the Korea Mouse Phenotyping Center using a Minispec LF50 (Bruker, Billerica, Massachusetts).

**Body temperature measurement**

Body temperature was measured using a rectal thermometer (testo 925, Testo SE, Titisee-Neustadt, Germany) by inserting a small-diameter temperature probe (length, 2 cm) into the anus of each mouse.

**Glucose tolerance test**

For the glucose tolerance test, the mice were fasted for 8 hours with access to water *ad libitum*. The mice were then injected with glucose (2 g/kg, i.p.). Glucose levels in tail nick blood were measured after 0, 15, 30, 60, and 120 minutes using a commercial glucometer.

**Immunostaining**

Cultured cells and cryo-sectioned brain slices were fixed with 4% PFA in PBS and blocked in 0.3% Triton X-100 with 5% normal goat serum for 1 hour. Then, they were incubated with primary antibodies (see Supplemental Table S1 for detailed information) overnight at 4°C. The secondary antibodies used for visualization were Cy3 (1:1000, Jackson Immunoresearch Laboratories, West Grove, Pennsylvania) and Alexa Fluor 488 (1:1000, Life Technologies, Carlsbad, California). The stained cells were mounted using VECTASHIELD with DAPI mounting solution (Vector Laboratories, Burlingame, California), and photographs were obtained using an epifluorescence microscope (Leica) or confocal microscope (Leica TCS SP5). For in vitro culture wells (coverslips), immunostained and DAPI-stained cells were counted in whole area or 5-8 random areas using an eyepiece grid at a magnification of 100× or 200×, and total positive cells (or percentages) in a well were calculated. For in vivo hypothalamic brain sections, immunoreactive cells were counted every 10 serial sections of the whole target region, and then the total number of cells was calculated by multiplication according to these factors. The Abercrombie correction factor [$N = n$ X $T/(T + D)$], where $N$ is the actual number of cells, $n$ is the number of nuclear profiles, $T$ is the section thickness (30μm), and $D$ is the average diameter of nuclei, was used to compensate for double counting in adjacent sections.

**Quantitative RT-PCR analysis**

Total RNA was prepared using TRIzol reagent (Thermo Fisher Scientific) according to the manufacturer's protocol, followed by cDNA synthesis using a SuperScript kit (Thermo Fisher Scientific). Quantitative RT-PCR was performed on a CFX96TM real-time system using iQ$^{TM}$ SYBR Green Supermix (Bio-Rad Laboratories, Hercules, California), and gene expression levels were determined relative to β-actin levels. Primer information is shown in Table S2.

**Western blotting**

Cell lysates were prepared in RIPA buffer (50 mM Tris-HCl pH 7.4, 150 mM NaCl, 1% Triton X-100, 1% sodium deoxycholate, 0.1% SDS, and 1 mM EDTA) containing a proteinase inhibitor cocktail and phosphatase inhibitor, followed by sonication. Lysates were cleared of insoluble material by centrifugation at 15,000 g and 4°C for 5 minutes, and the protein concentration was determined using a Pierce BCA assay (Thermo Fisher Scientific). SDS-PAGE was conducted followed by electrophoretic transfer to nitrocellulose membranes. Immunoblots were blocked with 5% bovine serum albumin (BSA) dissolved in tris buffer saline with 0.1% tween-20 (TBST) for 1 hour and then incubated with primary antibodies (see Supplemental Table S1 for detailed information) overnight at 4°C. For immunodetection of the primary antibodies, a goat-anti-rabbit-HRP conjugate or goat-anti-mouse-HRP conjugate (Cell Signaling Technology, Danvers, Massachusetts) was used at 1:1,000 in 5% BSA dissolved in TBST. HRP was detected using West-Pico chemiluminescence substrate (Thermo Fisher Scientific).

**Bone density measurement**

The tibia and femur from 16 m.o. mice were dissected and fixed overnight in 4% PFA in PBS. The bone was placed in scanning tubes (diameter, 12.3 mm) and imaged using a desktop µCT scanner (SkyScan 1076, Bruker) or Quantum GX Micro-CT Imaging System (PerkinElmer, Hopkinton, Massachusetts). The scans were integrated into 3D voxel images, and a region of interest (ROI) composed of 100 slices starting from 0.5 mm proximal to growth plate, constituting 1.6 mm in length, was chosen for analysis. The following 3D indices (BMD, BV/TV, Tb.N., Tb.Th.) were analyzed in the defined ROI.

**Bone histomorphometry**

The femur samples, which had been dissected and fixed, were decalcification in 20% EDTA (pH 7.4) at 4°C for a duration of 2 weeks and embedded in paraffin. Sections of the femur were stained with H&E to observe gross structure. Additionally, the samples were stained either for tartrate-resistant acid phosphatase (TRAP) to analyze osteoclasts or for alkaline phosphatase (ALP) to analyze osteoblasts using the TRAP/ALP stain kit (Wako Pure Chemical Industries, Tokyo, Japan). Briefly, the deparaffinized and rehydrated sections were incubated at 37°C for 30 minutes in each respective buffer. The slides were then mounted using VECTASHIELD (Vector Laboratories, Burlingame, California). Images were captured using the Nikon Digital Camera DXM1200 microscope system (Nikon Corporation, Tokyo, Japan). The TRAP-positive and ALP-positive results were normalized with respect to the bone surface.

**Serum proteome analysis**

Blood from 16 m.o. mouse was collected and allowed to clot for 1 hours at room temperature before centrifuging for 15 minutes at 2000 x g. Serum was collected and stored at -80 ℃. Proteome analysis was performed using a Proteome Profiler Mouse XL Cytokine Array (R&D Systems, Minneapolis, Minnesota) according to the manufacturer’s protocol.

**Serum leptin assay**

The leptin assay was conducted using a Mouse/Rat Leptin Quantikine ELISA Kit (R&D Systems) according to the manufacturer’s protocol.

**Bulk RNA-seq transcriptome analysis**

Total RNA was isolated from pelleted cells (~2 million cells) using TRIzol reagent. A cDNA library was prepared from high quality RNA (RIN >8) using TruSeq Stranded mRNA LT (Illumina, San Diego, California) and sequenced using a NovaSeq 6000 (Macrogen, Seoul, South Korea). The paired-end library was aligned to the UCSC hg19 human reference genome using the HISAT2 pipeline. DESeq2 was used for calling DEGs. The GTEx data used in this study were obtained from the dbGaP release, in which four random samples from the hypothalamus, hippocampus, cortex, and substantia nigra of a post-mortem donor were selected for comparison (study accession: phs000424.v7.p2; https://www.ncbi.nlm.nih.gov/projects/gap/cgi-bin/study.cgi?study_id=phs000424.v7.p2).

**Single-cell RNA-seq analysis**

Hypo-NSCs were cultured for 70 days in neuron differentiation medium as a neurosphere. Single-cell suspensions were obtained by enzymatic dissociation using papain solution (PAP2, Worthington Industries, Columbus, Ohio) and DNAse solution (D2, Worthington Industries) at a 20:1 ratio for 20 mins at 37°C. The cell suspensions were then centrifuged for 5 min at 500 g and resuspended in differentiation medium with 0.004% BSA. Cell viability was checked using an automated cell counter. Single-cell cDNA libraries were generated using a Chromium Next GEM Single Cell 3’ RNA library v3.1 reagent kit according to the manufacturer’s protocol (10x Genomics, Pleasanton, California). Single-cell libraries were sequenced on a NextSeq 500 platform to obtain 91- and 28-bp paired end reads. Cell Ranger (v. 5.0.0, 10x Genomics) was used to de-multiplex and quantify the unique molecular identifiers (UMIs). Acount matrix was generated for each sample with the default parameters, and the genes were mapped to the GRCh38 human reference genome.

The retrieved count matrix was then analyzed in the R (v. 4.1.1) environment. Doublets were identified and removed using R package scDblFinder (v. 1.10) [5]. Data processing was performed using the Seurat package (v. 4.0.3) [6]. Only cells with transcript numbers >2000, <40000, and <5% mitochondria transcripts were used for further processing. Counts were normalized using Seurat’s *SCTransform(vst.flavor = "v2")* function. The top 30 principal components (PCs) from *RunPCA()* were used for further analysis, following the standard Seurat workflow. Cell clustering was done using the Leiden algorithm. Cells were annotated manually based on well-known marker and automatically using ScType [7]. Voxel-based mapping was done using Voxhunt

(v. 1.01) [8]. The pseudo-time analysis was done using monocle3 (v. 1.2.9) [9]. For the regulon analysis, pySCENIC (v. 0.11.2) was run from a docker image on Jupyter Notebook using only protein-coding genes as input [10]. The regulons were then binarized based on k-means clustering, and regulon specificity was scored based on the Jensen-Shannon divergence. The UMAP embedding for neuron was examined using Sleepwalk (v. 0.3.2) [11]. For benchmarking with human fetal hypothalamus, integration was performed using CCA in the Seurat package from the SCT-transformed values, regressing against maternally imprinted gene *MEG3* to remove imprinting effects.

**Single-nucleus RNA-seq analysis**

Three mice per group were anesthetized and transcardially perfused with ice cold PBS made using DEPC-treated water. Entire hypothalami were dissected by isolating the brain parenchyma inside the circle of Willis. Non-hypothalamic tissue, including the optic tract, was removed, and then each sample was frozen at -80°C until nucleus isolation occurred. The frozen tissues were cut into small pieces and lysed with an EZ nucleus isolation kit (Sigma, NUC-101) containing RNase inhibitor (Promega). After Hoechst33342 nucleus-staining solution was added, pure nuclei were sorted using a FACS-ARIA II (BD). Each nucleus suspension was processed to generate single-nucleus RNA-seq libraries using the Chromium Next GEM Single Cell 3' RNA library v3.1 reagent kit according to the manufacturer's protocol (10x Genomics, Pleasanton, California), and sequencing libraries were sequenced on an Illumina HiSeq2500 according to the manufacturer's instructions (Illumina). Both reads were aligned to the Mm10 mouse genome reference sequence and quantified using Cell Ranger (v. 5.0.1, 10x Genomics). The UMI count matrix was then exported to R and analyzed using the Seurat package. We detected some batch effects in the form of significantly different distributions of ribosomal and mitochondrial transcripts. Therefore, the batch was used as a covariate during the downstream analyses. Droplet-based protocols such as this one suffers from extremely high dropout rates and sparse data due to technical problems [12]. Therefore, we performed data imputation using SAVER (v. 1.1.3) to estimate the missing values based on gene-to-gene relationships [13]. However, imputation protocols often lead to over-imputation and the loss of true 'biological zeroes,' which ultimately produces inaccurate results. To prevent that, we also performed scRecover (v. 1.12) to statistically estimate the probability of 'biological zeroes' and 'technical zeroes' [14]. Using moderate imputation, we were thus able to decrease the zero value in our dataset from 93.89% to 43.63% and increase the median detected genes from 1,183 to 4,665, which increased the detection power of the subsequent analyses. We confirmed that the imputation did not interfere with the clustering result. Clustering was performed in Seurat using the FindCluster function with the Leiden algorithm at a resolution =0.5. The RunUMAP function was used with the first 50 PCs for UMAP dimensional reduction. Marker gene expression was analyzed using FindAllMarkers() in Seurat using "scale.data" slot with logfc.treshold = 0.25, min.pct = 0.5, and only.pos = T. The ClusterProfiler (v. 4.4.4) package was used for the GO enrichment analysis [15]. Redundant ontologies were filtered using the simplify function with a 0.5 cutoff value. To compare data from aging control versus transplanted, DEGs were extracted from each cluster. Genes were considered significant if the adjusted p-value < 0.01 and the avg_diff > 0.5. The gene sets were then used as input for AddModuleScore. The cell-to-cell communication analysis was performed using CellChat (v. 1.4.0) [16]. ggplot2 and ggpubr were used to visualize the data, and stat compare means was used for post-hoc statistical analysis.

**Supplemental Table S1. List of reagents and resources used in this study**

| REAGENT or RESOURCE | SOURCE | IDENTIFIER |
|---|---|---|
| Antibodies | | |
| Rabbit polyclonal anti-Ki67 (1:1000) | Abcam | Cat# ab15580, RRID:AB_443209 |
| Mouse monoclonal anti-NKX2.1 (1:100) | Thermo Fisher Scientific | Cat# MA5-13961, RRID:AB_10984070 |
| Rabbit polyclonal anti-SOX2 (1:200) | Millipore | Cat# AB5603, RRID:AB_2286686 |
| Rabbit polyclonal anti-Rx (1:200) | Takara Bio | Cat# M228 |
| Mouse monoclonal anti-Nestin (1:500) | BioLegend | Cat# 656802, RRID:AB_2562474 |
| Mouse monoclonal anti-Islet 1 Homeobox (1:10) | DSHB | Cat# 40.2D6, RRID:AB_528315 |
| Mouse monoclonal anti-Tubulin beta 3 (TUBB3) (1:200) | BioLegend | Cat# 801202, RRID:AB_10063408 |
| Mouse monoclonal anti-MAP2 (1:200) | Sigma-Aldrich | Cat# M1406, RRID:AB_477171 |
| Mouse monoclonal anti-NeuN (RBFOX3) (1:200) | Millipore | Cat# MAB377, RRID:AB_2298772 |
| Rabbit polyclonal anti-Synapsin 1 (1:200) | Millipore | Cat# AB1543, RRID:AB_2200400 |
| Rabbit polyclonal anti-aMSH (1:1000) | Phoenix Pharmaceuticals | Cat# H-043-01, RRID:AB_10013604 |
| Rabbit polyclonal anti-Neuropeptide Y (1:1000) | ImmunoStar | Cat# 22940, RRID:AB_2307354 |
| Rabbit polyclonal anti-SCGN (1:500) | Sigma-Aldrich | Cat# HPA006641, RRID:AB_1079874 |
| Rabbit polyclonal anti-MCH (1:500) | Sigma-Aldrich | Cat# M8440, RRID:AB_260690 |
| Rabbit polyclonal anti-GFAP (1:500) | Agilent | Cat# Z0334, RRID:AB_10013382 |
| Mouse monoclonal anti-Vimentin (V9) (1:200) | Thermo Fisher Scientific | Cat# MA5-11883, RRID:AB_10985392 |

| | | |
|---|---|---|
| Rabbit polyclonal anti-Aquaporin 4 (249-323) (1:200) | Alomone Labs | Cat# AQP-004, RRID:AB_2039734 |
| Mouse monoclonal anti-A2B5, clone A2B5-105 (1:200) | Millipore | Cat# MAB312, RRID:AB_94709 |
| Rabbit polyclonal anti-phospho-STAT3 (Tyr705) (1:1000) | Cell Signaling Technology | Cat# 9131, RRID:AB_331586 |
| Rabbit polyclonal anti-STAT3 (1:1000) | Cell Signaling Technology | Cat# 9132, RRID:AB_331588 |
| Rabbit polyclonal anti-phospho-FoxO1 (Ser256) (1:1000) | Cell Signaling Technology | Cat# 9461, RRID:AB_329831 |
| Rabbit polyclonal anti-FoxO1 (C29H4) (1:1000) | Cell Signaling Technology | Cat# 2880, RRID:AB_2106495 |
| Mouse monoclonal anti-GAPDH (0411) (1:1000) | Santa Cruz Biotechnology | Cat# sc-47724, RRID:AB_627678 |
| Mouse monoclonal anti-NCAM (ERIC 1) (1:500) | Santa Cruz Biotechnology | Cat# sc-106, RRID:AB_627128 |
| Mouse monoclonal anti-GFAP (1:400) | BioLegend | Cat# 837201, RRID:AB_2565371 |
| Mouse monoclonal anti-Mitochondria (113-1) (1:500) | Abcam | Cat# ab92824, RRID:AB_10562769 |
| Rabbit polyclonal anti-NeuN (RBFOX3) (1:200) | Millipore | Cat# ABN78, RRID:AB_10807945 |
| Mouse monoclonal anti-MTCO1 (1D6E1A8) (1:1000) | Thermo Fisher Scientific | Cat# 459600, RRID:AB_2532240 |
| Mouse monoclonal anti-SDHB (G-10) (1:1000) | Santa Cruz Biotechnology | Cat# sc-271548, RRID:AB_10659104 |
| Rabbit polyclonal anti-FGF21 (1:1000) | Abcam | Cat# ab137715, RRID:AB_2833254 |
| Rabbit polyclonal anti-Beta Actin (1:1000) | Proteintech | Cat# 20536-1-AP, RRID:AB_10700003 |
| Rabbit monoclonal anti-phospho-eIF2alpha (Ser51) (119A11) (1:1000) | Cell Signaling Technology | Cat# 3597, RRID:AB_390740 |
| Rabbit monoclonal anti-eIF2alpha (D7D3) (1:1000) | Cell Signaling Technology | Cat# 5324, RRID:AB_10692650 |

| | | |
|---|---|---|
| Rabbit monoclonal anti-ATF-4 (D4B8) (1:1000) | Cell Signaling Technology | Cat# 11815, RRID:AB_2616025 |
| Goat polyclonal anti-GDF15 (1:1000) | Abcam | Cat# ab39999, RRID:AB_732535 |
| Mouse monoclonal anti-Monocytes / Macrophages, clone ED-1 (1:500) | Millipore | Cat# MAB1435, RRID:AB_177576 |
| Mouse monoclonal anti-Integrin alphaM (CD11b), clone OX-42 (1:500) | Millipore | Cat# CBL1512, RRID:AB_93253 |
| Rabbit polyclonal anti-Iba1 (1:500) | FUJIFILM Wako Shibayagi | Cat# 019-19741, RRID:AB_839504 |
| Rabbit monoclonal anti-Phospho-NF-κB p65 (Ser536) (93H1) (1:500) | Cell Signaling Technology | Cat# 3033, RRID:AB_331284 |
| Mouse monoclonal anti-TNF alpha, clone 52B83 (1:500) | Abcam | Cat# ab1793, RRID:AB_302615 |
| Mouse monoclonal anti-IL-1β (3A6) (1:500) | Cell Signaling Technology | Cat# 12242, RRID:AB_2715503 |
| Mouse monoclonal anti-Caspase-1 (p20) (1:500) | AdipoGen | Cat# AG-20B-0042-C100, RRID:AB_2755041 |
| Rabbit polyclonal anti-c-FOS (1:500) | Abcam | Cat# ab190289, RRID:AB_2737414 |
| Mouse monoclonal anti-YB-1 (E-7) (1:1000) | Santa Cruz Biotechnology | Cat# sc-398146 |
| Chemicals, Peptides, and Recombinant Proteins | | |
| L-Ascorbic acid | Sigma-Aldrich | Cat# A92902 |
| B-27™ Supplement (50X), minus vitamin A | Gibco | Cat# 12587010 |
| Poly-L-Ornithine | Sigma-Aldrich | Cat# P3655 |
| Fibronectin | Sigma-Aldrich | Cat# F1141 |
| SB431542 | Tocris | Cat# 1641 |
| ROCK inhibitor Y27632 | Millipore | Cat# 688000 |
| Purmorphamine | Millipore | Cat# 540223 |
| CHIR99021 | Stemgent | Cat# 04-0004-10 |
| IWP-2 | | |
| XAV | Sigma-Aldrich | Cat# X3004 |
| N6,2′-O-Dibutyryladenosine 3′,5′-cyclic (db-cAMP) | Sigma-Aldrich | Cat# D0260 |
| Recombinant human CNTF | Peprotech | Cat# 450-13 |
| Recombinant human/murine FGF-8b | Peprotech | Cat# 100-25 |
| Recombinant human noggin | Peprotech | Cat# 120-10C |
| Recombinant human sonic hedgehog (Shh) | Peprotech | Cat# 100-45 |
| Recombinant human insulin | Gibco | Cat# 12585-014 |
| Recombinant human FGF basic | R&D Systems | Cat# 233-FB |

| | | |
|---|---|---|
| Recombinant human/murine/rat BDNF | Peprotech | Cat# 450-02 |
| Recombinant human GDNF | Peprotech | Cat# 450-10 |
| Recombinant human leptin | Peprotech | Cat# 300-27 |
| Recombinant human ghrelin | Tocris | Cat# 1463 |
| Papain (PAP2) | Worthington | Cat# LK003176 |
| Deoxyribonuclease I (D2) | Worthington | Cat# LK003170 |
| D-(+)-Glucose | Sigma-Aldrich | Cat# |
| TRIzol™ Reagent | Invitrogen | Cat# 15596018 |
| 4% Paraformaldehyde Phosphate Buffer Solution | FUJIFILM Wako Shibayagi | Cat# 163-20145 |
| Critical Commercial Assays | | |
| LIVE/DEAD Viability/Cytotoxicity Kit | Invitrogen | Cat# L3224 |
| Pierce™ BCA Protein Assay Kit | Thermo Fisher Scientific | Cat# 23225 |
| SuperScript™ III Reverse Transcriptase | Invitrogen | Cat# 18080093 |
| iQ™ SYBR® Green Supermix | Bio-Rad | Cat# 1708880 |
| Proteome Profiler Mouse XL Cytokine Array | R&D Systems | Cat# ARY028 |
| Mouse/Rat Leptin Quantikine ELISA Kit | R&D Systems | Cat# MOB00B |
| Deposited Data | | |
| hPSCs-derived htNSC bulk RNA-seq | This paper | GSE214877 |
| hPSCs-derived htNSC organoid scRNA-seq | This paper | GSE215174 |
| htNSC-transplanted aged mouse hypothalamus snRNA-seq | This paper | GSE215174 |
| Experimental Models: Cell Lines | | |
| WAe009-A (H9) | WiCell | Cat# WA09, RRID: CVCL_9773 |
| CMC-hiPSC-0011 | National Stem Cell Bank (KSCB) | RRID:CVCL_WR33 |
| CS-001 | Corestem, Inc. | N/A |
| Experimental Models: Organisms/Strains | | |
| C57BL/6J mouse 12 months-old, male | Korea Basic Science Institute (KBSI) | N/A |
| C57BL/6J mouse 11 months-old, male | Jackson Laboratory | N/A |
| Oligonucleotides | | |
| See table S2 for primer list | This paper | N/A |
| Software and Algorithms | | |
| ImageJ (v. 1.53f51) | NIH | https://imagej.net/ImageJ |
| Prism 8 | GraphPad Software | https://www.graphpad.com/scientific-software/prism/ |

| | | |
|---|---|---|
| SPSS Statistics 26 | IBM | https://www.ibm.com/products/spss-statistics |
| ANY-maze | Stoelting | https://www.any-maze.com/ |
| Minispec Plus | Bruker | N/A |
| SkyScan 1076 | Bruker | N/A |
| Oxymax (v. 5.40.14) | Columbus Instrument | N/A |
| CLAX (v. 2.2.15) | Columbus Instrument | N/A |
| Leica Application Suite X | Leica | https://www.leica-microsystems.com/ |
| CaseViewer (v. 2.4) | 3DHISTECH Ltd. | https://www.3dhistech.com/solutions/caseviewer/ |
| CFX manager (v. 3.1) | Bio-Rad | https://www.bio-rad.com/ |
| Image Lab (v. 6.1) | Bio-Rad | https://www.bio-rad.com/ |
| R (v. 4.1.1) | R Core Team | https://www.r-project.org/ |
| Python (v. 3.8.14) | Python Software Foundation | https://www.python.org/ |
| Cell Ranger (v. 5.0.0 / v. 5.0.1) | 10x Genomics | https://www.10xgenomics.com/ |
| Seurat (v. 4.0.3) | [6] | https://satijalab.org/seurat/ |
| Voxhunt (v. 1.2.9) | [8] | https://quadbiolab.github.io/VoxHunt/ |
| Monocle3 (v. 1.0.1) | Cao et al., 2019 | https://cole-trapnell-lab.github.io/monocle3/ |
| pySCENIC (v. 0.11.2) | Aibar et al., 2017 | https://scenic.aertslab.org/ |
| scDblFinder (v. 1.10) | [5] | https://github.com/plger/scDblFinder |

| | | |
|---|---|---|
| SAVER (v. 1.1.3) | [13] | https://mohuangx.github.io/SAVER/ |
| scRecover (v. 1.12) | Miao et al., 2019 | https://miaozhun.github.io/scRecover/ |
| ClusterProfiler (v. 4.4.4) | Yu et al., 2012 | https://guangchuangyu.github.io/software/clusterProfiler/ |
| CellChat (v. 1.4.0) | Jin et al., 2021 | https://github.com/sqjin/CellChat |
| ScType | Ianevski et al., 2022 | https://github.com/IanevskiAleksandr/sc-type |
| Sleepwalk (v. 0.3.2) | Ovchinnikova et al. 2020 | https://anders-biostat.github.io/sleepwalk |
| Custom script | This paper | https://github.com/yasulistio/aging-hypothalamus |

424

**Supplemental Table S2. Primer list for qRT-PCR**

| Gene | Forward | Reverse |
|---|---|---|
| *NKX2-1* | CAGGACACCATGAGGAACAGCG | GCCATGTTCTTGCTCACGTCCC |
| *RAX* | GCAAGGTCAACCTACCAGAGGT | GCAGCTTCATGGAGGACACTTC |
| *OTP* | AGCTTCGCCAAGACTCACTACC | CACGGAACACGTTGGTCGTCTT |
| *ISL1* | GCAGAGTGACATAGATCAGCCTG | GCCTCAATAGGACTGGCTACCA |
| *SF-1* | CCAGACCTTCATCTCCATCGTG | TGGCGGTAGATGTGGTCGAACA |
| *EN1* | GCAACCCGGCTATCCTACTTATG | ATGTAGCGGTTTGCCTGGAAC |
| *YBX1* | TCGCAACGAAGGTTTTGGGA | TTCTTCTTTATGGCAGTCTGGTGT |
| *ASCL1* | TCTCATCCTACTCGTCGGACGA | CTGCTT CCAAAGTCCATTCGCAC |
| *ACTB* | GTGACAGCAGTCGGTTGGAG | AACAACGCATCTCATATTTGGAA |
| *Adrb1* | TTCTCCTAGAGGGCAAACCTTGT | CAGAGTGAGGTAGAGGACCCACA |
| *Adrb2* | CTGTGCCTTCGCAGGTCTTC | |
| *Adrb3* | CGACATGTTCCTCCACAAATCA | TGGATTCCTGCTCTCAAACTAACC |
| *Dio2* | TGGATTCCTGCTCTCAAACTAACC | CGCACACCAGTGAGCTCTGA |
| *Lipe* | GGGAGTGGGCTGTGCCTTA | TCCCCCAGGTTCAGCTTTT |
| *Actb* | CCTTCCAGCAGATGTGGATCA | CTCAGTAACAGTCCGCCTAGAA |
